# A streamlined tandem tip-based workflow for sensitive nanoscale phosphoproteomics

**DOI:** 10.1101/2022.04.12.488038

**Authors:** Chia-Feng Tsai, Yi-Ting Wang, Chuan-Chih Hsu, Reta Birhanu Kitata, Rosalie K. Chu, Marija Velickovic, Rui Zhao, Sarai M. Williams, William B. Chrisler, Marda L. Jorgensen, Ronald J. Moore, Ying Zhu, Karin D. Rodland, Richard D. Smith, Clive H. Wasserfall, Tujin Shi, Tao Liu

**Affiliations:** Biological Sciences Division, Pacific Northwest National Laboratory, Richland, WA 99354, USA; Institute of Plant and Microbial Biology, Academia Sinica, Taipei, Taiwan; Environmental Molecular Sciences Laboratory, Pacific Northwest National Laboratory, Richland, Washington 99354, USA; Department of Pathology, Immunology, and Laboratory Medicine, Diabetes Institute, College of Medicine, University of Florida, Gainesville, Florida 32611, USA

**Keywords:** Tip IMAC, Boosting to Amplify the Signal with Isobaric Labeling (BASIL), near single-cell, nanoscale phosphoproteome, Surfactant-assisted One-Pot (SOP) sample preparation, spatial phosphoproteome mapping

## Abstract

Effective phosphoproteome of nanoscale sample analysis remains a daunting task, primarily due to significant sample loss associated with non-specific surface adsorption during enrichment of low stoichiometric phosphopeptide. We developed a novel tandem tip phosphoproteomics sample preparation method that is capable of sample cleanup and enrichment without additional sample transfer, and its integration with our recently developed SOP (Surfactant-assisted One-Pot sample preparation) and iBASIL (improved Boosting to Amplify Signal with Isobaric Labeling) approaches provides a streamlined workflow enabling sensitive, high-throughput nanoscale phosphoproteome measurements. This approach significantly reduces both sample loss and processing time, allowing the identification of >3,000 (>9,500) phosphopeptides from 1 (10) µg of cell lysate using the label-free method without a spectral library. It also enabled precise quantification of ∼600 phosphopeptides from 100 cells sorted by FACS (single-cell level input for the enriched phosphopeptides) and ∼700 phosphopeptides from human spleen tissue voxels with a spatial resolution of 200 µm (equivalent to ∼100 cells) in a high-throughput manner. The new workflow opens avenues for phosphoproteome profiling of mass-limited samples at the low nanogram level.

## Introduction

The ability to trace dynamic protein phosphorylation in small populations of cells can reveal cell-to-cell heterogeneity in cell signaling (e.g., cellular responses to stimulus), and provide a foundation for identifying rare cell types within a clinical sample and better understanding of disease mechanisms. Conventional phosphoproteomic approaches require large sample sizes, which obscures cell type-specific signaling information. Sensitive and nanoscale phosphoproteomic methods have the potential to make such cell type-specific applications more feasible in clinical settings, and also enable more precise phenotypic analysis of cell (sub)types in a time- and/or dose-dependent fashion. Indeed, mass spectrometry (MS)-based single-cell proteomics has recently made significant advancements in detection sensitivity and high-throughput measurements^1-4^. A remaining technical challenge for nanoscale proteomics is the comprehensive quantitative profiling of post-translational modifications (PTMs), particularly protein phosphorylation states^5^, key indicators for signaling pathways and network activations essential for many physiological functions^6, 7^.

The combination of metal affinity chromatography (e.g., Fe(III) and TiO_2_) with extensive sample fractionation has made it possible to identify and quantify >30,000 phosphorylation sites from bulk materials (over mg of proteins)^8-10^. To extend phosphoproteomics measurement to small sample, Humphrey et al.^11^ described an EasyPhos protocol to facilitate rapid and reproducible phosphopeptide enrichment. They also modified the buffer during digestion and employed extended MS/MS accumulation times to enhance phosphopeptide detection in small-sized samples. Starting with 12.5 μg and 25 μg of peptides, approximately 8,000 and 9,000 phosphopeptides could be identified, respectively^11, 12^. Similarity, Post et al.^13^ assessed an automated enrichment protocol using Fe(III)-IMAC (immobilized metal affinity chromatography) cartridges on a AssayMAP Bravo platform to identify approximately 1,500 and 4,500 phosphopeptides from 1 μg and 10 μg of HeLa cell digests, respectively^13^. Chen et al.^14^ developed an integrated strategy, termed Phospho-SISPROT to identify approximately 600 and 3,000 phosphopeptides from 1 μg and 10 μg of HEK293T cell digests, respectively. However, technical challenge remains for effective phosphopeptide enrichment and deep phosphoproteome profiling from low µg and sub-μg sample sizes.

Recently, a signal ‘boosting’ strategy utilizing multiplexed isobaric labeling, in which a large amount of relevant ‘boosting’ (or ‘carrier’) material is labeled with one or several of the isobaric (tandem mass tag; TMT) reagent(s) and then mixed with the labeled low-amount samples using other TMT channels, has enabled single-cell proteomics analysis with enhanced detection sensitivity and sample throughput^15-17^. This strategy has been applied to enhance the detection of phosphopeptides in low amount of samples. It enabled the reliable quantification of >20,000 phosphorylation sites in 150 human pancreatic islets^18^ and >2,300 low abundant phosphotyrosine peptides from 1 mg of T cell receptor stimulated Jurkat T cells^19^.

To enable the sensitive phosphoproteome analysis of nanogram samples, herein we developed a streamlined tandem tip-based sample preparation workflow, which integrates Surfactant-assisted One-Pot (SOP) digestion, tandem tip-based C18-IMAC-C18 enrichment/cleanup, and improved Boosting to Amplify Signal with Isobaric Labeling (iBASIL) for MS data acquisition. The tandem tip method significantly reduces sample loss and increases throughput. Its analytical merits were benchmarked using 0.1, 1 and 10 ng peptides samples (equivalent to single cells, 10 cells, and 100 cells, respectively) from 3 different acute myeloid leukemia (AML) cell lines. The performance of the workflow was further demonstrated in the analysis of 100 Fluorescence-activated cell sorting (FACS)-sorted MCF10A cells before and after EGF treatment and laser-dissected human spleen tissue voxels (equivalent to ∼100 cells). This workflow can recapitulate the dynamic changes in protein phosphorylation in the small-sized samples, demonstrating its potential for broader applications in biological and biomedical research.

## Results

### The integrated tandem tip-based nanoscale phosphoproteomics workflow

Similar to single-cell proteomics^20^, the major challenge for nanoscale phosphoproteomics is the substantial sample loss due to non-specific surface adsorption during multi-step sample transfer steps. To address this issue, we have developed a streamlined ‘tandem tip’-based workflow for sensitive nanoscale phosphoproteomics (Figure 1) which effectively integrates four components: 1) a sample lysis and digestion by surfactant-assisted one-pot processing (SOP)^20^, 2) isobaric labeling/boosting using TMTpro (16-plex), 3) phosphopeptide enrichment/cleanup using a tandem C18-IMAC-C18 tip arrangement, and 4) improved MS data acquisition using the iBASIL strategy^17^. The streamlined nanoscale phosphoproteomic workflow primarily benefits from the new tandem tip C18-IMAC-C18 method that maximizes phosphopeptide recovery from IMAC enrichment of low-input samples.

**Figure 1.**
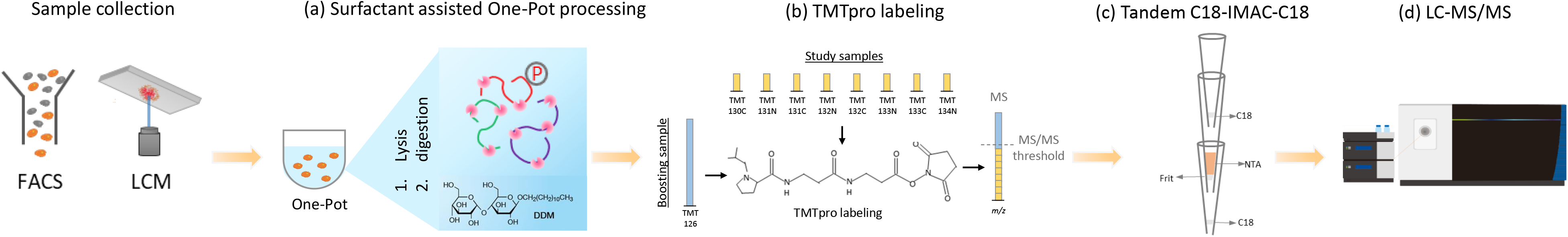
A streamlined tandem tip-based workflow for nanoscale phosphoproteomics. (a) The proteins are digested by our recently developed SOP (Surfactant-assisted One-Pot) approach. (b) The digested peptides are labeled with TMTpro reagents for sample multiplexing. (c) TMT labeled phosphopeptides are purified by tandem tip-based C18-IMAC-C18. (d) The enriched phosphopeptides are analyzed by LC-MS/MS applying iBASIL settings.

Conventional workflows for phosphopeptide enrichment have generally been developed for bulk sample analysis with typical ≥100 µg sample inputs^10^. They are usually performed in a sample vial (i.e., in-vial processing)^10^, involve multiple sample transfers, use large processing volumes and incur long processing times. Our alternative tandem tip approach integrated C18 tip-based peptides desalting with IMAC-based phosphopeptide enrichment followed by direct LC-MS analysis without sample transfer steps. The Ni-NTA silica beads were packed into the pipet tip to form tip IMAC. The C18 elution buffer for the desalting procedure (80% ACN in 0.1% TFA) is also used as the IMAC loading buffer. The IMAC elution buffer (NH_4_H_2_PO_2_ at pH 4.4) is compatible with reversed-phase C18 for sample loading, which eliminates the step of peptide lyophilizing. The buffer compatibility makes the tandem C18-IMAC-C18 method ideal for phosphopeptide enrichment from small samples and subsequent LC-MS/MS analysis.

### Tip-IMAC vs. in-vial IMAC

Compared to in-vial phosphopeptide enrichment, tip-IMAC reduced the total processing time from ≥100 min to ∼20 min (Figure 2a) and thus increased the analytical throughput by 5X. The performance of tip-IMAC step was evaluated using both unlabeled and TMT-labeled peptides. The same amount of MCF-7 peptides was aliquoted for tip-IMAC and in-vial IMAC^10^. When comparing the tip-IMAC protocol with in-vial IMAC protocol^10^ (Figure 2b), we observed 22% (from 10,490 to 12,769) and 12% (from 7,590 to 8,504) increases in the numbers of phosphopeptide identification for samples without and with TMTzero labeling, respectively. Besides the increased number of phosphopeptides, the tip-IMAC method resulted in higher specificity (>90%) of phosphopeptide purification for both without and with TMT labeling when compared to in-vial IMAC (Figure 2c).

**Figure 2.**
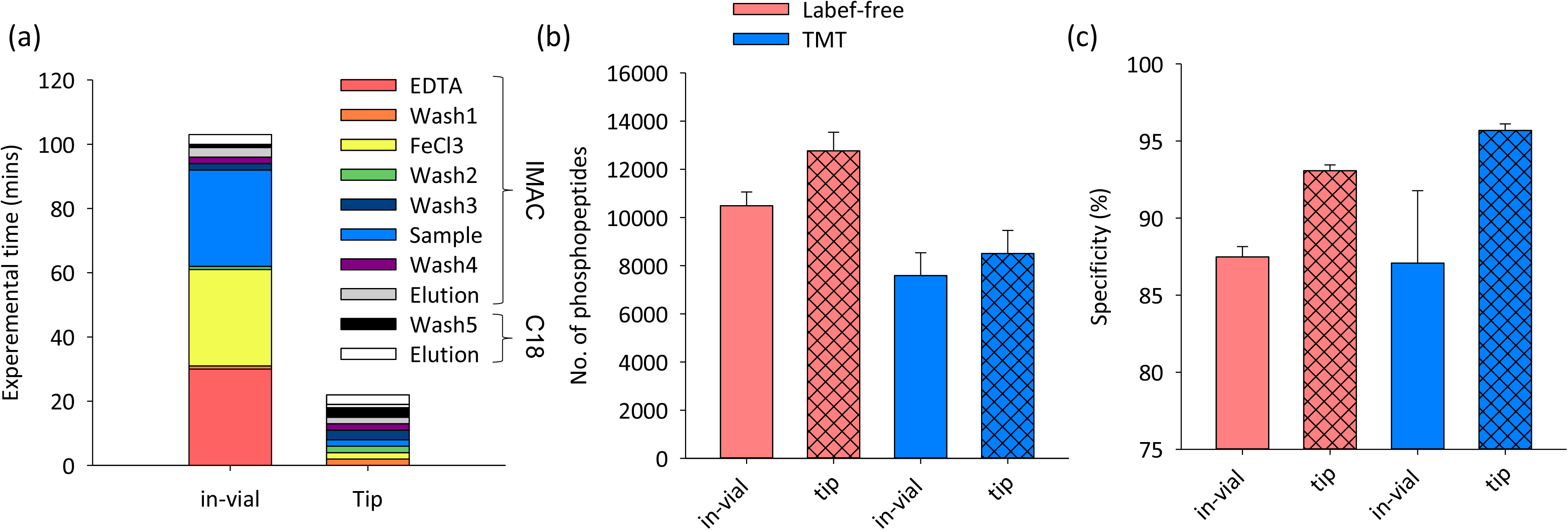
Comparison of in-vial and tip-based IMAC for phosphopeptide enrichment. (a) The processing time for in-vial and tip-based IMAC. The number of identified phosphopeptides (b) and the specificity of phosphopeptide enrichment (c) for unlabeled and TMT-labeled peptides from MCF-7 cells using the in-vial and tip-based IMAC.

### Tandem tip vs. step-by-step enrichment

We integrated an integrated tandem tip-based method (C18-IMAC-C18) to avoid sample transfer between different vials and further increase sample recovery (Figure S1a). The digested peptides from 10 µg of MCF-7 cell lysates were purified in parallel by the tandem tip and conventional (‘step-by-step’) methods. Compared to the step-by-step method, the tandem tip method improved 1.04-fold in the number of identified phosphopeptides (from 9,557 to 9,955) and 1.03-fold in the XIC area (Figure S1b). Moreover, this tandem tip method also provided improved Pearson correlation with lower quantitation CV (coefficient of variation) (Figure S1c). In addition, we evaluated the inter-day and intra-day reproducibility of this integrated C18-IMAC-C18 method using more precise SRM (Selective Reaction Monitoring)-based targeted proteomics approach. We spiked 10 SIL phosphopeptide mixture at 2 µM concentration into large amount of reference endometrial tumor digest (200 µg) for tandem tip purification. Its high reproducibility across different batches and days was evident by the Pearson correlation (r>0.99) and CV (median=15.8%) (Figure S2).

### Tandem IMAC-HpH-C18 for phosphopeptide fractionation

The home-made tip IMAC can be directly integrated with a high-pH C18 reversed-phase (HpH-C18) tip^21^ for in-tip phosphopeptide fractionation. Purified phosphopeptides from 20 µg peptides were directly fractionated by HpH-RP tip (Figure S3a) into 4 fractions. After LC-MS/MS analysis, the higher separation efficiency (0.9 % overlapping phosphopeptides across 4 fractions) by HpH-RP tip Figure S3b) effectively reduced sample complexity and increased the number of identified phosphopeptides (n=15,906) compared to single-shot LC-MS/MS (n=9,214).

The integrated IMAC-HpH-RP tip can be also used for large sample size (e.g., ≥100 µg) phosphopeptide fractionation. The phosphopeptides from 500 µg peptides were firstly purified by tip-IMAC and directly fractionated into 6 fractions by high-pH reversed-phase (HpH-C18) tip^21^ (Figure S3d). The strong fractionation capability of this method (Figure S3e) is evident by the low overlap of identified phosphopeptides between fractions and the overall comprehensive phosphoproteome coverage (46,703 identified phosphopeptides and 6,095 phosphoproteins) from only 500 µg peptides. This is especially useful for generating spectral libraries for highly effective DDA (Data-dependent acquisition) and DIA (Data-independent acquisition) analysis (see Figure 3 below). These results demonstrated that tandem C18-IMAC-C18 can be broadly used for sensitive phosphoproteomic analysis of small samples as well as a convenient fractionation system (without the need of LC) to achieve in-depth phosphoproteome coverage.

**Figure 3.**
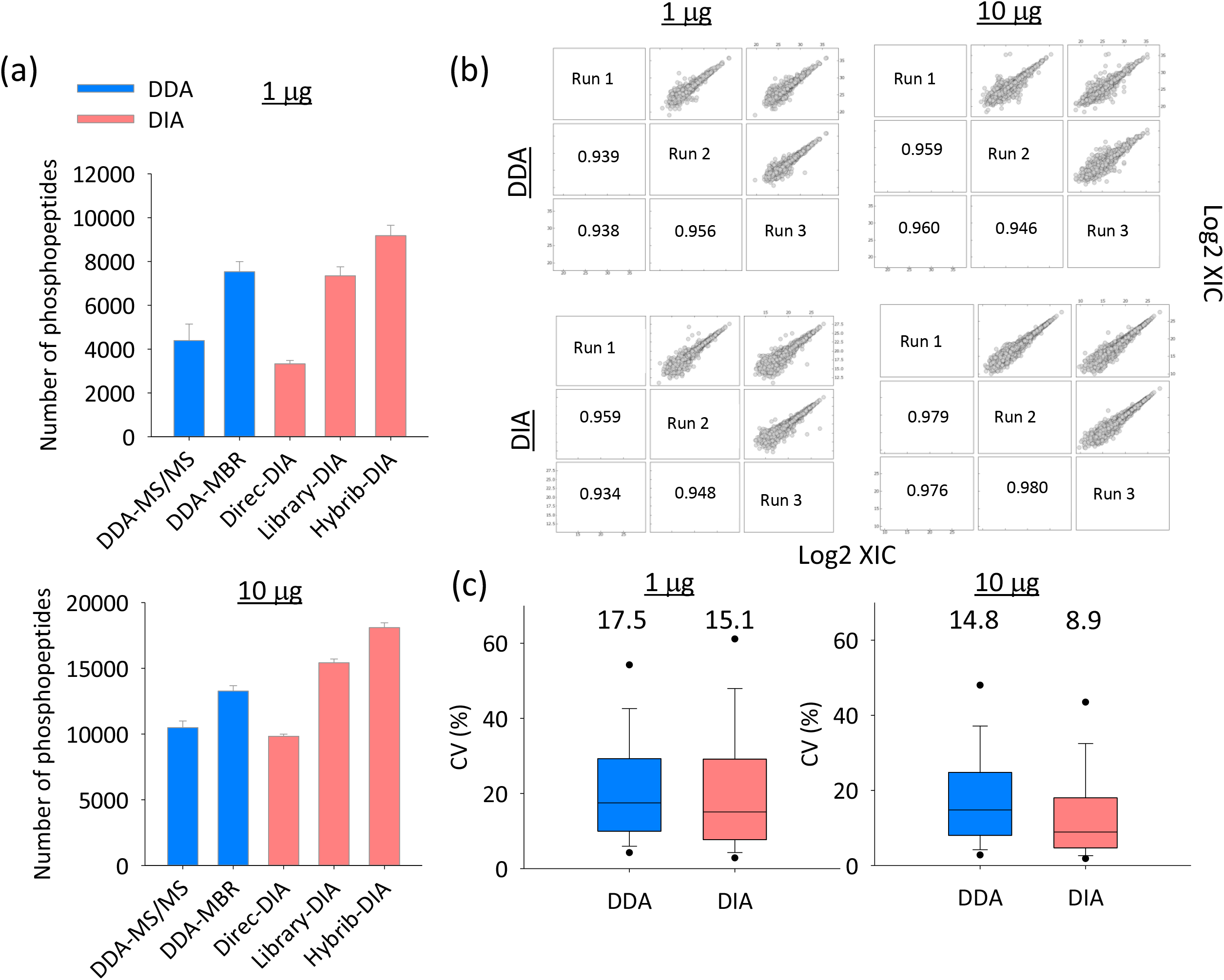
Evaluation of the performance of tandem C18-IMAC-C18 using DDA and DIA. The number of phosphopeptides (a), reproducibility (Pearson correlation) (b) and CV (%) (c) of phosphoproteomics analyses of 1 and 10 µg of proteins from mixture of 3 AML cells using single-shot DDA and DIA. A phosphoproteome spectrum library constructed from data described in Figure S3d and S3e was used for Library DIA and MBR DDA analysis.

### Evaluation of tandem C18-IMAC-C18 detection sensitivity using label-free methods

We first evaluated the detection sensitivity of tandem tip using two different label-free methods (DIA and DDA). The tip-IMAC allowed for identification of 3,330 (9,831) and 4,394 (10,482) phosphopeptides in 1 µg (10 µg) of tryptic peptides by single-shot direct DIA and DDA without library search or MBR (MS/MS only), respectively (Figure 3a). To improve the phosphoproteome coverage, we used the phosphopeptides identified from the above HpH-C18 tip-fractionated 500- µg sample (Figure S3d and S3e) to construct a spectral library. This enabled a substantial increase of phosphopeptide identifications: 9,180 (18,100) and 7,535 (13,268) phosphopeptides in 1 µg (10 µg) peptide samples by DIA (library search) and DDA (MBR), respectively (Figure 3a); DIA also was demonstrated with better quantitation quality as indicated by low protein CVs (Figure 3b and 3c).

We next evaluated the detection sensitivity of the tandem C18-IMAC-C18 method using DDA analysis of A549 with 1-, 10-, and 50-µg protein input (Figure 4a), as well as an even lower input of 0.1-µg (equivalent to 1,000 cells). The tandem tip method allowed for identification of 12,975 and 9,675 phosphopeptides from 50 and 10 µg of starting material, respectively, and 3,412 phosphopeptides from 1 µg of starting material. After using the MBR function across different samples, the number of identified phosphopeptides was increased to 4,477, 12,824 and 15,754 from 1, 10, and 50 µg, respectively.

**Figure 4.**
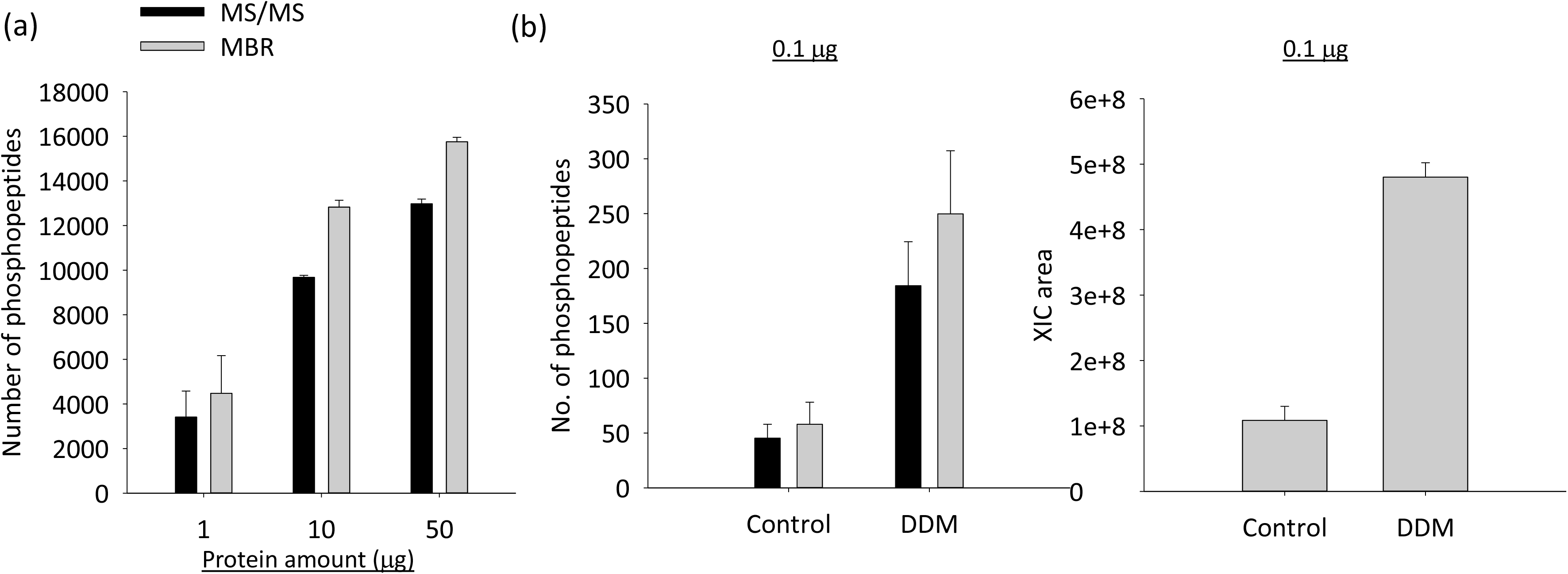
Detection sensitivity of tandem C18-IMAC-C18. (a) The numbers of identified phosphopeptides (with and without MBR) from 1, 10 and 50 µg of A549 cell lysates. (b) The numbers of identified phosphopeptides (with and without MBR) and XIC area for 0.1 µg of cell lysates with and without DDM.

We also assessed the effect of DDM (Figure 4b), which has been previously demonstrated to greatly reduce surface adsorption loss in single-cell and low-input proteomics^20, 22, 23^, for enhancing phosphopeptide detection at sub-µg input level using the tandem tip method. 0.1 µg of cell lysates were digested in the PCR tubes without and with DDM treatment (i.e., the SOP approach^20^), and loaded into tandem tip (C18-IMAC-C18) for phosphopeptide enrichment with and without DDM treatment. With the surface protection by DDM, the number of identified phosphopeptides was substantially increased from 45 to 184 based on the MS/MS (or 58 to 250 based on MBR), with ∼4.4-fold enhancement in MS signals (XIC areas) (Figure 4b). In summary, we demonstrated that the tandem tip method enables robust and comprehensive phosphoproteome coverage for ≥1 µg of proteins. However, the coverage is still limited for sub- µg input levels even when spectrum library is used for matching, presumably due to the lower MS1 signals.

### Integration of tandem C18-IMAC-C18 method with iBASIL

To further increase the detection sensitivity for sub-µg input levels, we integrated the tandem C18-IMAC-C18 method with our recently developed carrier proteome-based iBASIL approach^17, 18^ (Figure 1d). To mimic nanoscale phosphoproteome analysis, 0.1, 1 and 10 ng of tryptic digest of 3 different AML cells (MOLM-14, K562 and CMK) equivalent to total protein content in 1, 10 and 100 cells, respectively, were analyzed in 3 separate TMT-16 experiments, each with 1 µg of the same tryptic digest of mixed cells as the boosting sample (Figure 5a). The 3 sets of TMT16-labeled samples were then processed by tandem tip. To minimize the ion undersampling effect caused by the overwhelming amount of ions from the boosting channel^17, 24^, high AGC (1E6) and long ion injection time (IT, 0.5 s and 1.5 s) were used to improve the sampling of phosphopeptide ions in sample channels. Approximately 2,800 phosphopeptides were identified for all three TMT-16 experiments, but only 36, 290 and 2,103 phosphopeptides were quantified (70% valid values in one cell type) from the 1-, 10- and 100-cell analysis using the 500-ms IT time setting, respectively (Figure 5b). Increasing the IT time from 0.5 s to 1.5 s increased the cycle time, resulting in decreasing the number of identified phosphopeptides down to ∼1,700; however, higher IT time effectively improved the ion sampling from the samples in the sample channel, resulting in significantly higher number of quantified phosphopeptides (e.g., from 290 to 928 for the 10-cell analysis) (Figure 5b). Higher IT time slightly decreased the number of quantifiable phosphopeptides for the 100-cell (10 ng) analysis (Figure 5b), presumably due to the overall reduced number of PSMs as a result of the prolonged duty cycle. Wider dynamic range of TMT reporter ion intensity was obtained by increasing the IT time (Figure 5c). The higher IT time also improved the quantitation precision in CV (%) (Figure 5e) and separation of the three AML cell types in 100-cell analyses using the resulting phosphoproteome profiles (Figure 5f). Moreover, the higher quality quantitative results (Figure 5g) via higher IT time generated more statistically significant phosphopeptides amongst the three types of AML cells (618 and 1,099 phosphopeptides with adjusted p value <0.05 using the 0.5-s and 1.5-s methods, respectively), resulting in increased coverage in signaling pathways (Figure S4).

**Figure 5.**
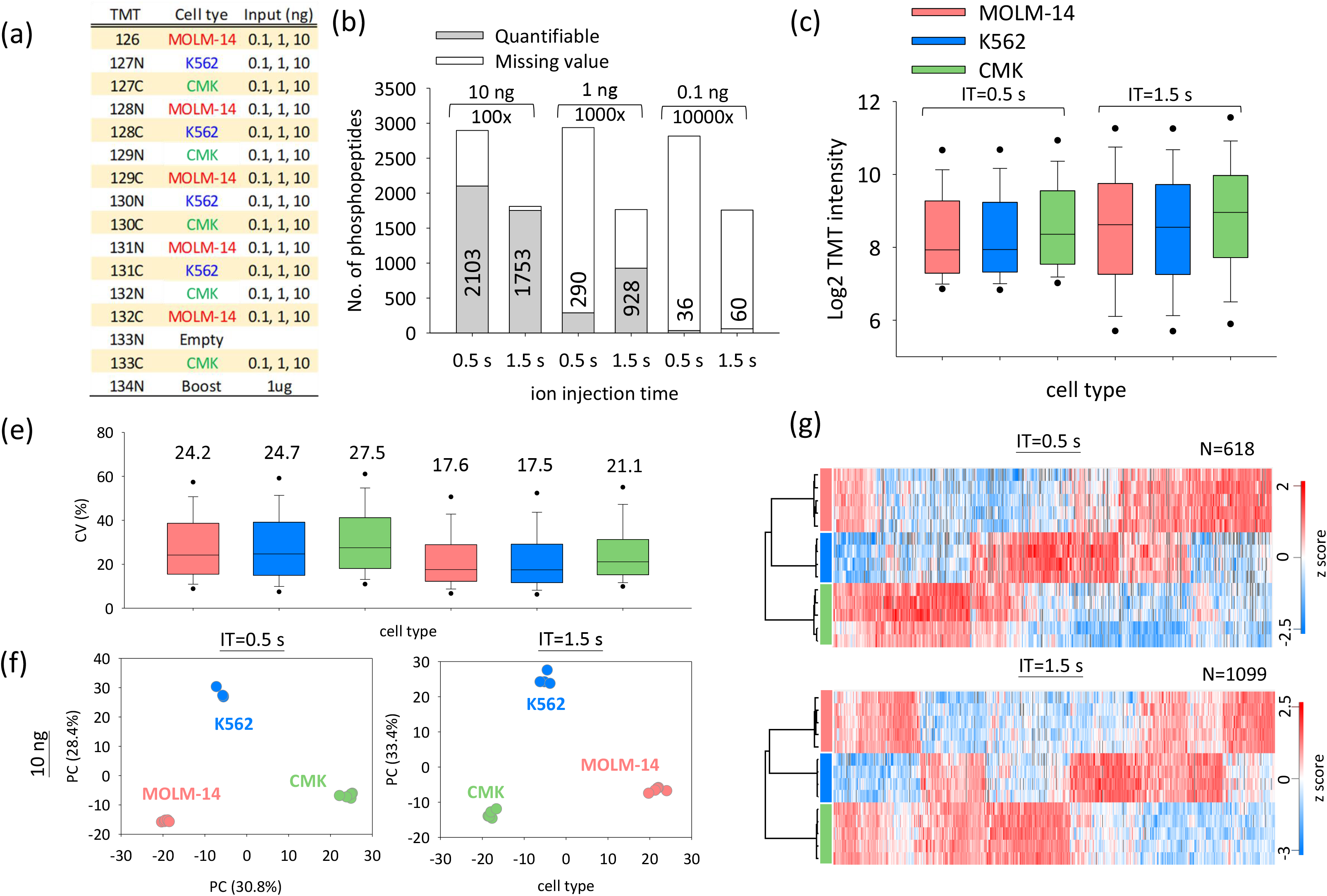
Mimic nanoscale phosphoproteome analysis using tip-IMAC and iBASIL. (a) The TMT experiment design for the phosphoproteomics analysis of tryptic digests (0.1, 1 and 10 ng) from three different types of AML cells (MOLM-14, K562 and CMK). (b) The numbers of quantified phosphopeptides (70% no-missing value in study samples) from 0.1, 1 and 10 ng of tryptic digests under different ion injection times (0.5 and 1.5 s). The TMT reporter ion intensity distribution (c), CV (%) of replicate experiments (e), PCA analysis (f), and heatmap of significantly changed phosphopeptides of the 10-ng input samples under different ion injection times (g).

Up to 928 phosphopeptides can be quantified in the 10-cell analysis (1 ng of peptides); however, the three AML cell types did not cluster tightly in the principal component analysis (PCA), especially when lower IT time was used (Figure S5). The outlier data points were from three TMT sample channels (TMT132N, 132C and 133C), most likely due to isotopic impurities from the TMT134N channel where the boosting material was labeled at 1,000-fold higher input level. Similar result was observed in previous study where TMT-10/11/16 was used^17, 25^. This suggests that more channels adjacent to the boosting channel should be left empty when a high boosting ratio is used (e.g., ≥1,000x). Nevertheless, these results demonstrated the utility of the integrated C18-IMAC-C18/iBASIL platform for robust quantitative nanoscale phosphoproteomic analysis of 10-100 cells.

### Streamlined SOP/C18-IMAC-C18/iBASIL workflow for phosphoproteomics analysis of 100 sorted cells

To explore the full potential of the streamlined nanoscale phosphoproteomics platform, small numbers of cells sorted by FACS were used to evaluate its performance. 100 FACS-sorted MCF10A cells with and without EGF treatment were processed by SOP method, followed by 16-plexed TMTpro labeling (130C to 134N). To avoid the potential isotopic ‘leakage’ issue^17, 24^ (e.g., in the 10-cell analysis in Figure S5), the TMT-labeled 100-cell samples were mixed with 1 µg of pre-digested MCF10A tryptic peptides that was labeled in a far separate channel (TMT126) while leaving the TMT channels 127N to 130N empty (Figure 6a). The labeled peptides were directly loaded into tandem tip C18-IMAC-C18 for phosphopeptide enrichment/cleanup, and the resulting samples were analyzed by LC-MS/MS with the iBASIL settings (i.e., both high AGC and IT time).

**Figure 6.**
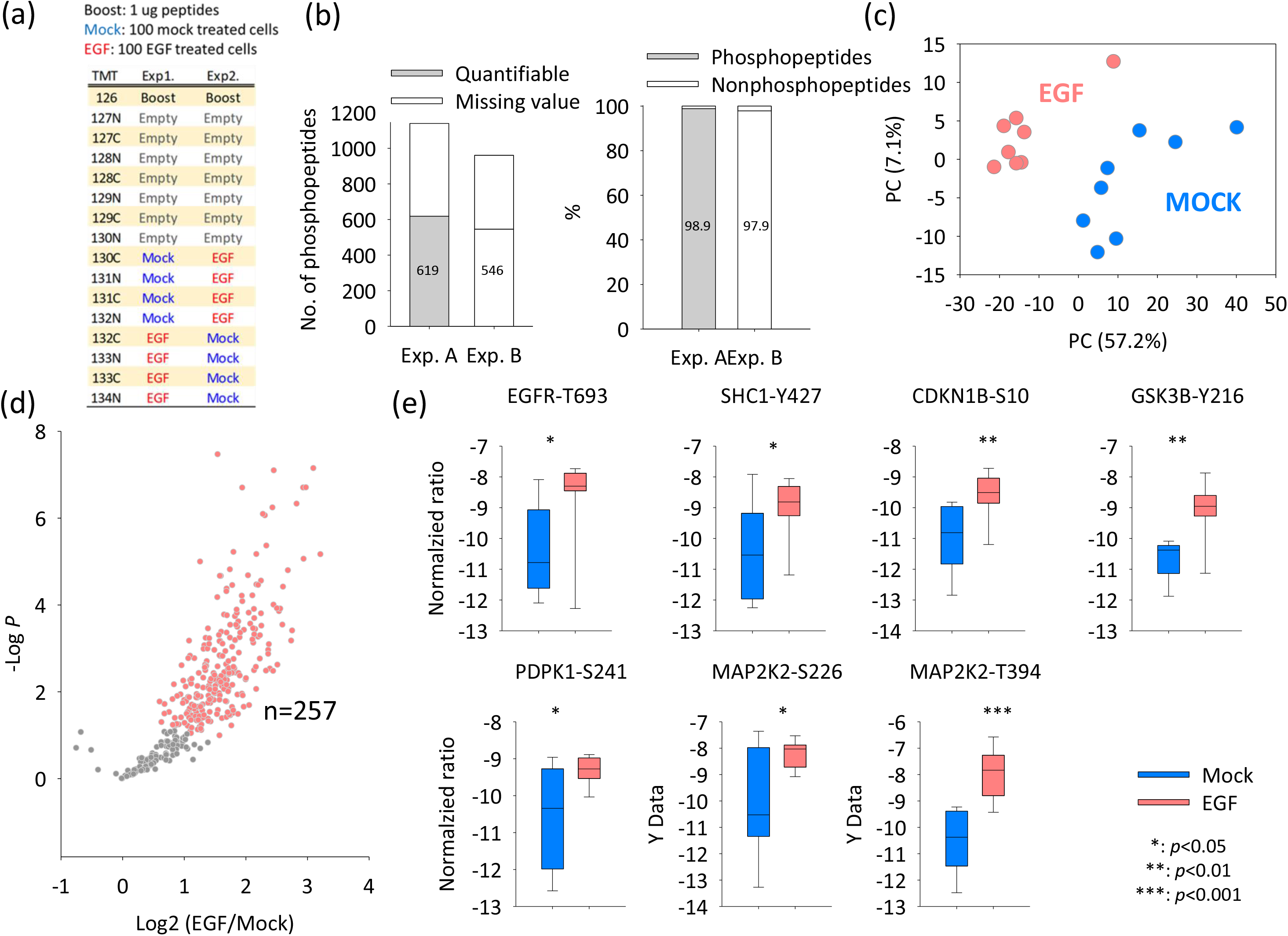
Phosphoproteome analysis of 100 MCF10A cells sorted by FACS using the streamlined SOP/C18-IMAC-C18/iBASIL workflow. (a) The TMT experiment design. (b) The number of quantified phosphopeptides (70% no-missing value in study samples) and the enrichment specificity in each TMT experiment. (c) PCA analysis shows the cells cluster under the two different conditions. (d) Volcano plot shows significantly changed phosphopeptides between EGF-treated cells and mock cells. (e) The significantly altered phosphorylation sites in the ErbB signaling pathway. Y-axis means the normalized TMT intensity after medium centering and batch correction by Combat.

Approximately 1,000 phosphopeptides were identified with ∼600 quantifiable sites in >70% of the study samples using half of the resulting sample with the conventional LC-MS platform (i.e., the Lumos Orbitrap MS and 75-µm i.d. LC column). High specificity (>98%) was achieved for phosphopeptide enrichment (Figure 6b), which helped reduce potential dynamic range compression due to co-isolated non-phosphopeptides during MS/MS sampling. The quantified phosphopeptides allowed for robust separation of 100 sorted MCF10A cells (single-cell level input of the enriched phosphopeptides) with and without EGF treatment (Figure 6c) and 257 phosphopeptides were found to be upregulated upon EGF treatment (Figure 6d and Table S2). The altered phosphopeptides were subjected to KEGG and Reactome pathway analysis using DAVID^26^ and STRING^27^, respectively. As expected, known pathways (e.g., ErbB and insulin signaling pathways; Table S2) were enriched. We identified 7 key upregulated phosphorylation sites in ErbB signaling pathways, such as T693 in EGFR^28^, Y427 in SHC1^29^, S10 in CDKN1B^30^, Y216 in GSK3B^31^, S241 in PDPK1^32^, and S226^33^ and T394 in MAP2K2^34, 35^ (Figure 6e). In addition, mRNA splicing was found to be activated after EGF treatment (Table S2). In a previous study, Zhou et al.^36^ described that EGF signaling promotes changes in splicing via AKT and SRPKs, which is consistent with our finding in this study. We also tried to analyze 10 sorted MCF10A cells with and without EGF treatment (Figure S6). While the different cell states (i.e., with and without EGF treatment) can be separated by PCA analysis, the coverage of quantifiable phosphopeptides were limited (n<200; enrichment specificity at >98% level), likely due to the insufficient detection sensitivity (Figure S6).

### Spatial phosphoproteomics and proteomics analysis using the streamlined workflow

We next tested the utility of the streamlined workflow for spatial phosphoproteomics analysis using small human spleen tissue voxels (Figure 7). Two distinct regions, red and white pulps, were dissected by laser capture microdissection (LCM) to generate small tissue voxels (Figure 7a). Each tissue voxel has a size of 200 µm × 200 µm with 10 µm in thickness (approximately 100 cells^22^]. They were first processed by convenient microPOTS (microdroplet processing in one pot for trace samples) at the low µL processing volume^37^. The streamlined C18-IMAC-C18/iBASIL workflow was then applied for nanoscale phosphoproteomics analysis.

**Figure 7.**
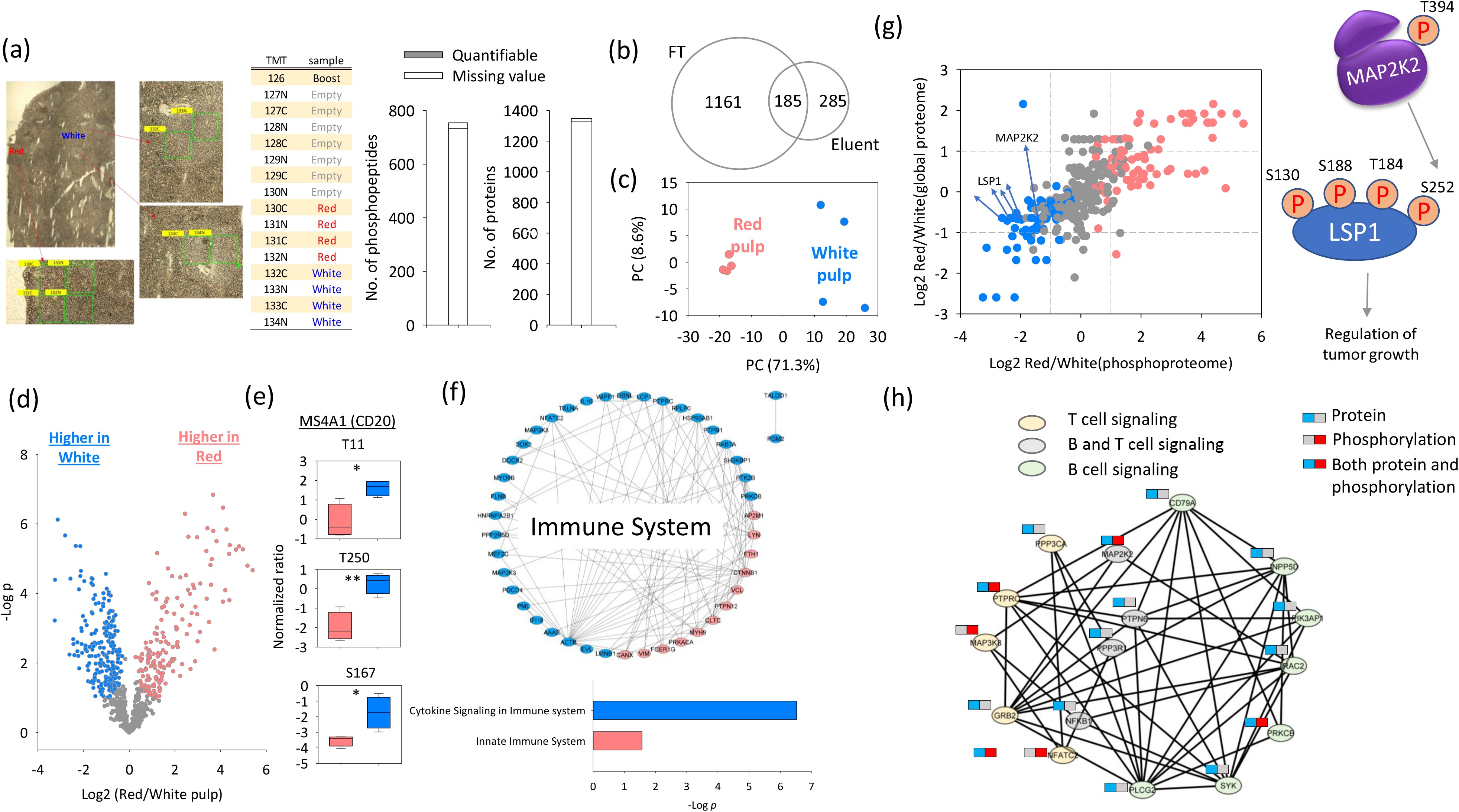
Phosphoproteome analysis of LCM-dissected human spleen tissue voxels. (a) The optical image of the white and red pulp regions, the TMT experiment design and the number of quantified phosphopeptides from IMAC eluent and proteins from IMAC flow-through (FT). (b) The overlap of quantified proteins between IMAC eluent and FT. (c) PCA of the phosphoproteome data. (d) Volcano plot shows significantly changed phosphopeptides in white pulp and red pulp. (e) The significantly altered phosphorylation sites in the MSFA1 (CD20) proteins. (f) The annotated pathway for altered phosphorylation events. (g) The quantitation correlation between global and phosphoproteome. (h) The annotated B- and T-cell signaling pathways for overexpressed phosphorylation peptides and proteins in white pulp.

Importantly, with TMT labeling and analysis of both the IMAC-enriched fraction and IMAC flow-through, we can characterize the phosphoproteome and proteome of each tissue voxel in an integrated fashion. A total of 737 unique phosphopeptides (470 phosphoproteins) were identified with 84% specificity from IMAC eluent, and 1,346 proteins (6,216 peptides) were identified from the IMAC flow-through (Figure 7a). Compared to the global proteome from the flow-through fraction, an additional 285 less abundant proteins were identified through IMAC enrichment (Figure 7b), such as the G protein-coupled seven transmembrane receptor CXCR5 and the transcription factor TFAP4. The red and white pulp regions could be readily separated in PCA using either the phosphoproteome data (Figure 7c) or the proteome data (Figure S7a). Analysis of variance (ANOVA) revealed that 56% of the quantified phosphopeptides (Figure 7d; FDR < 0.05) and 69% of the quantified proteins (Figure S7b; FDR < 0.05) were differentially expressed between the white and red pulp regions, including 11 CD markers (Figure S7c).

Red pulp is a loose spongy tissue with chords of reticular cells located between venous sinuses that primarily contain erythrocytes, with lymphocytes, macrophages, granulocytes, and plasma cells potentially trafficking through. Uniquely in the red pulp of human spleen CD8a lines these sinuses, termed littoral cells^38^, and also marks a subset of T cells through the spleen (CD8a red in Figure S7d). CD3e marks both CD8 and CD4 T cells interacting in follicles and dispersed throughout the organ (CD3e white in Figure S7d). This is the major site for both erythrocyte storage and removal of bound immune complexes through complement receptors. In addition, senescent erythrocytes are destroyed by macrophages with hemoglobin:haptoglobin complexes being loaded onto CD163 positive macrophages for metabolic processing. This is consistent with the observed expression of surface protein markers (e.g., CD99, CD47 and CD55) as well as other protein markers (e.g., haptoglobin and CD163) at higher levels in red pulp, with the platelet-related pathway enriched (Table S3).

In contrast, white pulp is composed of lymphoid follicles with central B lymphocytes (CD20 green in Figure S7d), interacting CD3 T cells and follicular dendritic cells orchestrating immune responses. There is also a collection of lymphocytes surrounding small arterioles (CD31 blue in Figure S8), called periarteriolar lymphoid sheaths (PALS)^39^. CD20 is a B cell-specific membrane protein and represents an attractive target for therapeutic antibodies^40^. As expected, CD20 protein (Figure S7c) and its phosphorylation site (Figure 7e) have higher expression levels in the white pulp. Both white and red pulps contain lymphocytes, neutrophils, macrophages, and other innate lymphoid cells^39^. Corresponding pathways relevant to these cell types were observed to be enriched in both regions of spleen tissues (e.g., immune system; Table S3 and Figure 7f). However, cytokine signaling was enriched in white pulp, reflecting the role of T cell derived cytokines in triggering maturation of B cells in white pulp, while proteins related to the innate immune system were enriched in red pulp, reflecting ongoing immune reactions in red pulp (Figure 7f). Interestingly, both LSP1 and an upstream kinase MAP2K2 had significant increase at both protein and phosphorylation site levels in the white pulp (Figure 7g). LSP1 is a key switch that triggers apoptosis in activated B cells^41^; increased MAP2K2 phosphorylation is consistent with the increased proliferation of immature B cells in response to the cytokines released by T cells in the white pulp.

Taken together, our results demonstrate unprecedented data quality in integrated, high-resolution spatial phosphoproteomics and proteomics analysis using the streamlined nanoscale proteomics workflow within an exemplar spleen organ. This approach has the potential to be an enabling tool for spatial phosphoproteomics profiling. These results also recapitulated at the protein level the cell type distribution and function of white pulp and red pulp, confirming the ability of spatially-resolved proteomics to capture important functional differences.

## Discussion

In this study, we developed and demonstrated a streamlined tandem tip-based workflow for sensitive and scalable nanoscale phosphoproteomics analysis that also integrated a front-end SOP for initial sample processing with minimal sample loss and the boosting/carrier-assisted iBASIL approach for multiplexed, sensitive, and effective MS analysis of the resulting nanoscale phosphoproteomics sample. This workflow was shown to recapitulate the major signaling pathway changes in as few as 100 mock and EGF-treated MCF10A cells and 200 µm × 200 µm × 10 µm human spleen tissue voxels.

Our home-made tandem tip-based method is a sensitive, robust and readily adoptable method for both label-free (DDA and DIA) and TMT-based nanoscale phosphoproteomics. The tandem tips can also be inserted into either Eppendorf vials or 96-well plates (via an adapter) for high-throughput phosphopeptide enrichment using centrifugation. We carefully evaluated the performance of this tandem tip-based method (e.g., sensitivity, reproducibility, and quantitation accuracy) using label-free MS analysis and low-input levels of proteins from cultured cells, demonstrating significantly improved results compared to previously reported methods/platforms. For label-free DDA analysis, the tandem tip-based method can identify 9,675 (12,824 after MBR) phosphopeptides from 10 µg of proteins from A549 cells. Using DIA, the number of phosphopeptides increases to 9,831 and 18,100 by direct- and library-DIA, respectively. Compared to previous studies (Table S4), the coverage is much higher than that from the optimized AssayMAP Bravo platform using Fe^3+^-IMAC cartridges (4,541 phosphopeptides without MBR from 10 µg of proteins from HeLa cells)^13^ and EasyPhos method using TiO_2_ (∼4,000 (∼8,000 after MBR) phosphopeptides from 12.5 µg of proteins from U-87 MG cells)^12^. Chen et al.^14^ further developed an integrated approach (termed Phospho-SISPROT) which used SCX/C18 tip for digestion and then transferred the digested peptides into Tip-IMAC/C18 tip. However, the phosphopeptides in the elution buffer can’t be directly loaded into C18 tip for desalting due to elution buffer contains higher organic solve (50% ACN) and higher pH salt (10%NH_3_·H_2_O). Therefore, only approximately 3,000 phosphopeptides can be identified from 10 μg of HEK293T cell digests after MBR (Table S4), respectively. This suggests that our tandem tip-based method reduces sample loss and provides higher phosphopeptide recovery (although the use of different LC and MS instruments and cell lines may still affect the final phosphoproteome coverage).

The same trend remains when the starting materials decreased to 1 µg or lower input levels (Table S4); however, the phosphoproteome coverage became much more limited, with only ∼200 phosphopeptides detected at the 0.1-µg input level by either our tandem tip method or the AssayMAP Bravo platform^13^. This is likely due to the lack of sufficient MS1 (precursor) ion intensities for triggering MS/MS in the current Orbitrap instruments. Indeed, although the IT accumulation time was set to 50 ms for MS1 data acquisition from 0.1 µg of cell lysate digests, the median of the actual IT time was only 4.7 ms (Figure S8). This is because uninformative singly charged ions dominated the MS spectra during full m/z range acquisition in the C-trap^42-44^. This issue can be alleviated by recent developments for improving the MS1 signal in label-free analysis. For example, Meier et al. reported a BoxCar approach which was built on the ability of the quadrupole–Orbitrap mass analyzer to be filled sequentially with ions in different mass windows to increase the actual ion injection time at MS1 level^43^. High field asymmetric ion mobility spectrometry (FAIMS)^45^ is another attractive option where an asymmetric electric field was used to selectively filter ion populations by varying the compensation voltage. FAIMS has been used to selectively remove +1 ions while broadly transmitting multiply charged peptides^3, 46^. Recently, Woo *et al*. developed an ion mobility-enhanced MS acquisition and peptide identification method, TIFF (Transferring Identification based on FAIMS Filtering), which significantly extends the ion accumulation times for peptide ions by filtering out singly charged background ions^44^. Note that a tradeoff from using high ion injection times to increase precursor ion sampling efficiency is that the cycle time is also increased significantly, which leads to substantially reduced number of MS/MS spectra for peptide identification. Therefore, the overall effectiveness of either BoxCar or TIFF approach mainly depends on matching MS1 features to an existing library via the MBR algorithm^47^. Nevertheless, integration of our tandem tip-based sample processing and enrichment method with the TIFF or BoxCar approach is still expected to increase the detection sensitivity for the label-free analysis of nanoscale phosphopeptides.

In contrast to the label-free analysis, the TMT-based boosting strategy can increase not only sample throughput (up to 18 channels)^48^ but also detection sensitivity with the use of the carrier proteome in the boosting channel for significantly improved MS1 signal. This boosting strategy has recently been demonstrated for effective single-cell proteomics analysis^1, 15, 17^. It has also been applied to study low abundant PTMs such as phosphotyrosine peptides^19^ or secreted glycoproteins^49^. When integrated with our streamlined tandem tip-based method, the boosting strategy enabled precise quantification of ∼600 phosphopeptides from 100 sorted cells (Figure 6) as well as >1,500 phosphopeptides from 10 ng of cell lysate digests (∼100 cells; Figure 5). To increase the ion sampling from the actual low-input sample under the masking by the boosting material^24^ [, we applied similar iBASIL settings^17^, i.e., high AGC (1E6) and high ion injection times (0.5 or 1.5 s in Figure 5 and 3 s in Figure 6). To minimize the impact of TMT reagent isotopic impurities on quantitation quality^50^, especially when higher boosting ratio is used (Figure S5), more TMT channels adjacent to the boosting channel can be kept empty (Figure 6). In addition, the use of narrower isolation windows (0.7 Da), and the high IMAC enrichment specificity achievable for TMT-labeled phosphopeptides (∼98% in Figure 6; ∼84% in Figure 7) using the tandem tip-based method, can also effectively reduce the ‘compression’ caused by the co-isolated and often relatively abundant non-phosphopeptides. Application of these approaches led to high ion purity (close to 1) for the detected phosphopeptides in 100 MCF10A cells and 100 cell-equivalent human spleen tissue voxels (Figure S9). Additional developments, such as the use of infrared photoactivation to boost reporter ion yield in isobaric tagging^51^, are expected to further improve the performance of iBASIL-like analysis.

To better realize the potential for single-cell PTM analysis, it has been suggested that the PTM peptides enriched from bulk samples may be mixed with the un-enriched single cells for direct LC-MS/MS analysis of the PTMs^52^. This strategy may reduce the sample loss for the single-cell samples, however, the co-isolation of phosphopeptides and non-phosphopeptides in MS/MS will continue to be a major issue for quantitation (reduction in quantitation values), in addition to the masking effect by the boosting material. Indeed, phosphoproteins are often of low abundance and phosphorylation events are commonly happening at low stoichiometry^53-55^. Furthermore, the physicochemical properties of phosphopeptides critically determine their ionization efficiency^56, 57^. For example, removing the phosphate group can increase the ion generation efficiency ∼10-fold compared to that of the original phosphopeptides^58^, which could greatly decrease the sensitivity for detection of the signal from the single-cell phosphopeptides, especially when the single-cell phosphopeptides and non-phosphopeptides are co-isolated for MS analysis. Therefore, to achieve highly effective single-cell phosphoproteome analysis, phosphopeptide enrichment is needed and the detection sensitivity needs to be further improved. One future development will focus on improvements in LC-MS detection sensitivity by effective integration of ultralow-flow LC and a higher efficiency ion source/ion transmission interface with the most advanced MS platform and data acquisition methods. Another direction is to utilize higher resolution IMS (e.g., structures for lossless ion manipulations, SLIM^59^ for gas-phase separation to reduce sample complexity and effectively remove unwanted ions (e.g., singly charged ions, co-isolated species and isotopic impurity), resulting in increased detection sensitivity and improved quantitation accuracy. To improve the robustness and throughput, the PTM enrichment processing can also implement automated small-volume liquid handling via the use of a 96-well plate adapter/holder.

## Conclusion

The ability to comprehensively characterize PTMs at a nanoscale or single-cell level is critical for better understanding of biological variability at functional levels. To this end, we have developed a readily implemented streamlined tandem tip-based method for nanoscale phosphoproteomics. This method capitalizes on seamless integration of C18 with IMAC to form tandem C18-IMAC-C18 for rapid, effective phosphopeptide enrichment. Further integration with our SOP and iBASIL methods forms a streamlined workflow allowing precise quantification of ∼600 phosphopeptides from 100 MCF10A cells sorted by FACS and ∼700 phosphopeptides from 200 µm × 200 µm × 10 µm human spleen tissue voxels (equivalent to ∼100 cells). Moreover, the resulting high-quality phosphoproteome data were able to capture important functional differences in the samples. This streamlined workflow should open new avenues for nanoscale phosphoproteome profiling of small numbers of cells and mass-limited samples at the low nanogram levels and high-resolution spatial phosphoproteome mapping of human tissues, which cannot be accessed by current proteomics platforms. With further improvements in detection sensitivity as well as sample processing, it has the potential to moving towards single-cell phosphoproteomics.

## Method

### Reagents

Urea, dithiothreitol (DTT), iodoacetamide (IAA), iron chloride, triethylammonium bicarbonate (TEABC), ethylenediaminetetraacetic acid (EDTA), ammonia phosphate (NH_4_H_2_PO_4_), trifluoroacetic acid (TFA), formic acid (FA), acetonitrile (ACN), n-Dodecyl β-D-maltoside (DDM), protease, phosphatase inhibitor, DMSO (HPLC grade), and Phosphate-Buffered Saline (PBS) were obtained from Sigma (St. Louis, MO). The Ni-NTA silica beads and agarose beads were obtained from Qiagen (Hilden, Germany) and the Empore™ extraction C18 disks were obtained from 3M (St. Paul, MN). TMT-16 reagents were purchased from Thermo Fisher Scientific (Waltham, MA). Water was processed using a Millipore Milli-Q system (Bedford, MA). Polypropylene microwell chip with 2.2-mm wells diameter was manufactured on polypropylene substrates by Protolabs (Maple Plain, MN). Tris(2-carboxyethyl) phosphine hydrochloride (TCEP-HCl), and 50% Hydroxylamine (HA) were purchased from Thermo Fisher Scientific (Waltham, MA). Ethanol was purchased from Decon Labs, Inc (King of Prussia, PA).

### Cell culture and protein digestion

The AML (MOLM-14, K562 and CMK), MCF-7, A549 and MCF10A breast cell lines were obtained from the American Type Culture Collection and were prepared as previously described^60^. Cell pellets were washed with ice-cold phosphate-buffered saline (PBS), lysed in a lysis buffer containing 50 mM TEABC, pH 8.0, 8 M urea, and a 1% protease and phosphatase inhibitor. The protein concentration was determined using the BCA protein assay (Thermo Fisher Scientific). The protein solutions were denatured with 10 mM DTT for 15 mins at 37 °C and alkylated with 50 mM iodoacetamide in the dark for 30 mins at room temperature. The resulting samples were diluted by 8-fold with 50 mM TEABC and digested by lysyl endopeptidase and trypsin (protein:enzyme, 20:1, w/w) at 37 °C overnight. The digested tryptic peptides were acidified by TFA with a final TFA concentration of 0.5%, and then desalted by C18 SPE extraction and concentrated for BCA assay.

### TMT labeling for the bulk samples

After digestion, the digested peptides (in 200 mM HEPES) were mixed with TMT-16 reagents dissolved in 100% acetonitrile (ACN) and allowed to react for 1 h. An optimized ratio of TMT to peptide amount of 1:1 (w/w)^61^ was used (i.e., 100 µg of peptides labeled by 100 µg of TMT reagent). The labeling reactions were stopped by adding 5% hydroxylamine (final concentration is 0.5%) for 15 min and then acidified with trifluoroacetic acid (TFA; final concentration is 0.5%). Peptides labeled with different TMT tags were mixed in the same tube, after which the ACN concentration was adjusted to below 5% (v/v) and the samples were desalted by C18 SPE.

### FACS sorting and protein digestion

For procedure of cell sorting has been descripted in the previous study^20^. The sorted cells were denatured with 1 µL 0.2%DDM with 10 mM TCEP in 50mM TEABC. Samples were incubated at 75 °C for 1 h for denaturation and reduction. After that, the samples were alkylated with 50 mM iodoacetamide in the dark for 30 mins at room temperature. The resulting samples were digested by 10 ng lysyl endopeptidase and 10ng trypsin at 37 °C overnight (The final volume was 3 µL). The digested tryptic peptides were diluted with 1 µL 200mM HEPES (pH= 8.8) labeled with 1 µL TMTpro reagent (20 µg/µL) for 1 hr at room temperature. Then, the labeling peptides were treated with 0.25 µL 5% HA for 15 mins and acidified by TFA. %). Peptides labeled with different TMT tags were mixed in the same tube, after which the ACN concentration was adjusted to below 5% (v/v) and the samples were loaded into tandem tip C18-IMAC-C18 for phosphopeptide enrichment/cleanup.

### Stable isotope-labeled peptides

Ten crude stable isotope-labeled (SIL) phosphopeptides were synthesized with 13C/15N on C-terminal lysine or arginine (New England Peptide, Gardner, MA). The SIL peptides were dissolved in 15% ACN and 0.1% FA at a concentration of 1.5 mM individually. A mixture of SIL peptides was made with a final concentration of 4 pmol/μL for each peptide.

### LC-MS/MS Analysis

For most of experiments, lyophilized peptides were reconstituted in 12 μL 0.1% TFA with 2% ACN containing 0.01% DDM^20^ and 5 μL of the resulting sample was analyzed by LC-MS/MS using an Orbitrap Fusion Lumos Tribrid mass spectrometer (Thermo Scientific) connected to a nanoACQUITY UPLC system (Waters) (buffer A: 0.1% FA with 3% ACN and buffer B: 0.1% FA in 90% ACN) as previously described^17^. Peptides were separated by a gradient mixture with an analytical column (75 μm i.d. × 20 cm) packed using 1.9-μm ReproSil C18 and with a column heater set at 50 °C. Peptides were separated by an LC gradient: 2-6% buffer B in 1 min, 6-30% buffer B in 84 min, 30-60% buffer B in 9 min, 60-90% buffer B in 1 min, and finally 90% buffer B for 5 min at 200 nL/min. For experiments in Figure 3, lyophilized peptides were reconstituted in 0.1% TFA with 2% ACN containing 0.01% DDM and injected using a PAL autosampler (CTC Analytics AG, Switzerland). The sample was concentrated into an online SPE column (150 μm i.d., 360 μm o.d., 4 cm long) and separated using a 50 μm i.d., 360 μm o.d., 50 cm long column packed with 3-μm C18 particles (300-Å pore size; Phenomenex, Terrence, CA). The nanoLC separation used a Dionex UltiMate NCP-3200RS system (Thermo Scientific, Waltham, MA) with mobile phases of water with 0.1% FA (buffer A) and ACN with 0.1% FA (buffer B). Peptides were separated through a linear gradient from 8% to 35% buffer B over 100 min at a flow rate of 150 nL/min. The separated peptides were analyzed using a Thermo Scientific Q Exactive Plus. Top 10 precursor ions were selected for MS/MS. Different AGC settings and maximum ITs at MS and MS/MS level for different DDA experiments were listed in Table S1. The DIA-MS/MS scan was performed in the HCD mode with the following parameters: precursor ions from 350–1650 *m/z* were scanned at 120,000 resolution with an ion injection time of 60 ms and an AGC target of 3E6. The scan rage of m/z (isolation window) of DIA windows from 377 (54), 419(32), 448(28), 473.5(25), 497.5(25), 520.5 (23), 542.5 (23), 564.5 (23), 587 (24), 610.5 (25), 635 (26), 660 (26), 685.5 (27), 712.5 (29), 741 (30), 771 (32), 803.5 (35), 838.5 (37), 877 (42), 921 (48), 972 (52), 1034.5 (71), 1133.5 (129) and 1423.5 (453) were scanned at 15,000 resolution with an ion injection time of 120 ms and an AGC target of 3E6. The isolated ions were fragmented with HCD at 30% level.

### SRM assay development and LC-SRM

To evaluate the peptide quality and select the best transitions for each peptide, heavy peptide mixtures were subjected to an initial analysis using a TSQ Altis triple Quadrupole Mass Spectrometer (Thermo Fisher Scientific) equipped with a nanoACQUITY UPLC system (Waters, Milford, MA). The collision energies for individual transitions were obtained by using empirical equations from the Skyline software. The 6 most intensive fragment ions for each peptide were initially selected. Next, the LC characteristics, MS response, interferences, and endogenous detectability of the SIL peptides were further evaluated by spiking them into water and pooled samples for IMAC enrichment and LC-SRM analysis. In the end, 3 or more transitions per peptide were selected for configuration of the final assays for reproducible targeted quantification. The eluent from IMAC was dissolved in 10 µL 0.1% TFA with 2% ACN containing 0.01% DDM and 9 μL of the resulting sample was loaded onto the LC column (100 µm i.d. × 10 cm, BEH 1.7-µm C18 capillary column (Waters)) and separated under the following conditions: mobile phases (A) 0.1% FA in water and (B) 0.1% FA in ACN; flow rate at 400 nL/min; 42-min gradient (min:%B): 11:0.5, 13: 5, 18:20, 23:25, 29:40, 31:95, 36:0.5. The LC column was operated at a temperature of 44 °C. The parameters of the triple quadruple instrument were set as follows: 0.7 fwhm Q1 and Q3 resolution, and 1 s cycle time. Data were acquired in the unscheduled SRM without blinding the samples.

### Data analysis for DDA experiment

The raw MS/MS data were processed with MSFragger via Fragpipe^62, 63^. The MS/MS spectra were searched against a human UniProt database (fasta file dated July 31, 2021 with 40,840 sequences which contain 20,420 decoys) and (initial) fragment mass tolerances were set to 20 ppm. A peptide search was performed with full tryptic digestion (Trypsin) and allowed a maximum of two missed cleavages. Carbamidomethyl (C) was set as a fixed modification; acetylation (protein N-term), oxidation (M) and Phospho (STY) were set as variable modifications for global proteome analysis. For match-between-run analysis, 10 ppm m/z tolerance, 0.4 mins RT tolerance and 0.05 MBR ions FDR were used for analysis. The final reports were then generated (PSM, ion, peptide, and protein) and filtered at each level (1% protein FDR plus 1% PSM/ion/peptide-level FDR for each corresponding PSM.tsv, ion.tsv, and peptide.tsv files). The intensities of all ten TMT reporter ions were extracted from Fragpipe outputs and analyzed by Perseus^64^ for statistical analyses.

### Data analysis for DIA experiment

The DIA data analysis was performed essentially as described in the DDA data analysis (above), except that a library file was further imported into DIA-NN^65^ for DIA library matching.

### Data analysis for SRM experiment

SRM data were analyzed using Skyline software (version 4.2). The total peak area ratios of endogenous light peptides and their heavy isotope-labeled internal standards (i.e., L/H peak area ratios) were calculated for quantification. Peak detection and integration were carried out based on two criteria: (1) same retention time and (2) similar relative SRM peak intensity ratios across multiple transitions between light (endogenous) peptides and the heavy SIL peptide standards. All data were manually inspected to ensure correct peak detection and accurate integration.

### CODEX Staining and Imaging

Barcoded antibody staining of tissue sections mounted on cover slips was performed using a commercially available CODEX Staining Kit according to the manufacturer’s instructions. Raw images were processed and stitched using CODEX Processor software including cycle alignment, drift compensation, background subtraction, and cell segmentation. Image analysis was performed with the Akoya Multiplex Analysis Viewer (MAV) in Fiji with KNN/FLANN clustering, gating and spatial network mapping.

## Data availability

The RAW global MS data and the identified results from Fragpipe have been deposited in Japan ProteOme STandard Repository (jPOST: https://repository.jpostdb.org/)^66^. The accession codes: JPST001468 for jPOST and PXD032019 for ProteomeXchange. The access link is https://repository.jpostdb.org/preview/189170729262298fbf5daee and access key is 4830. SRM results including the assay characterization data are organized as Skyline files on the Panorama server^67^ and can be accessed via https://panoramaweb.org/nanoscale_phosphoproteomics.url. Here are the reviewer account details; Password:oHgxvRzH. The index of all the source data was listed in Table S1.

## Acknowledgments

Portions of the research was supported by a UG3CA256967 grant (to TS) from the National Institutes of Health (NIH) Common Fund, Human Biomolecular Atlas Program (HuBMAP) grant, a U24CA210955 grant (to TL and RDS) and a U01CA214116 grant (to KDR) from the NCI Clinical Proteomic Tumor Analysis Consortium (CPTAC), a NCI Early Detection Research Network (EDRN) Interagency Agreement ACN20007-001 (to TL), a RF1MH128885 grant (to TS) from the Brain Initiative Cell Census Network (BICCN), a R01GM139858 grant (to TS) from the National Institute of General Medical Sciences (NIGMS), a R21CA223715 grant (to TS) from the NCI Innovative Molecular Analysis Technologies (IMAT), and a P41GM103493 grant (to RDS) from the National Institute of General Medical Sciences. The research was performed at the Environmental Molecular Sciences Laboratory (grid.436923.9), a Department of Energy Office of Science User Facility sponsored by the Office of Biological and Environmental Research and located at the Pacific Northwest National Laboratory.

## Author contributions

C.-F.T., T.L. and T.S. conceived and designed the study. C.-F.T. and Y.-T.W. performed all proteomics experiments and data analysis. W.B.C. conducted FACS experiments for cell isolation. C.H.W. and M.L.J. provided spleen tissue sample. S.M.W. and Y.Z. produced microchips. M.V. performed LCM collection and microPOTS digestion. R.J.M., R.Z. and R.K.C. maintained the performance of LC-MS/MS. C.-F.T., T.L., T.S., Y.-T.W., C.-C.H. and R.B.K. provided input on the experimental design, data presentation, and manuscript preparation. C.-F.T., T.L., T.S., Y.-T.W., D.S.W., H.L., C.H.W., R.D.S. and K.D.R. wrote the manuscript with input from all other authors.

## Corresponding authors

Correspondence to Tao Liu, Tujin Shi or Chia-Feng Tsai.

## Figure Legend

**Figure S1. The comparison of the step-by-step and integrated tip-based IMAC methods for phosphopeptides enrichment**. (a) Schematic illustration of the two methods. The number, XIC area (b), Pearson correlation, and CV (%) (c) of identified phosphopeptides purified from 10 µg of proteins from the A549 cell lysate.

**Figure S2. The reproducibility of integrated tip-IMAC for phosphopeptides enrichment**. Pearson correlation and CV (%) of peak areas of 10 spiked SIL phosphopeptides (SRM data) in a reference endometrial tumor sample enriched with tip-based IMAC in different days and batches.

**Figure S3. The IMAC-HpH-RP tip method for microscale phosphopeptide fractionation**. (a) Using the integrated C18-IMAC-HpH-RP tip method for phosphopeptides fractionation (4 fractions for 20 µg tryptic peptides of MCF-7 cells). (b) Low overlap of the phosphopeptides identified from the 4 different fractions. (c) The number of identified phosphopeptides using the single-shot and HpH-RP tip-based fractionation method. (d) The tandem IMAC-HpH-C18 tip for phosphopeptide fractionation (6 fractions for 500 µg of tryptic peptides of MCF-7 cells). (e) Separation efficiency calculated by the percentage of common phosphopeptides between fractions.

**Figure S4. Pathway enrichment analysis of significantly changed phosphopeptides of 10 ng tryptic digests of AML cells under different ion injection times**.

**Figure S5. Effect of ion injection time on quantitation quality in the mimic nanoscale phosphoproteome analysis**. The PCA analysis of quantified phosphopeptides of 10 ng tryptic digests of the AML cells under two different ion injection times (0.5 and 1.5 s).

**Figure S6. Phosphoproteome analysis for 10 sorted MCF10A cells**. (a) The numbers of quantified (70% no-missing value in study samples) phosphopeptides and enrichment specificity in each TMT experiment. (b) PCA analysis shows the clustering of cells from the two different treatment conditions.

**Figure S7. Global proteome analysis of LCM-dissected human spleen tissue voxels**. (c) PCA of the proteome data. (d) Volcano plot shows significantly changed proteins in white pulp and red pulp. (c) The altered proteins expression of surface markers. (d) Representative CODEX image of human spleen tissue: CD8α (red), CD163 (yellow), CD3e (white), CD20 (green), CD31 (blue). Scale bar, 560 µm.

**Figure S8. The ion injection time distribution of purified phosphopeptides from 0.1** µ**g proteins from the A549 cell lysate at MS1 (a) and MS2 level**.

**Figure S9. The ion purity distribution of detected phosphopeptides in Figure 6 and 7**.

**Table S1. The raw files and corresponding samples, experimental conditions and MS instrument setting**.

**Table S2. Pathway enrichment analysis of significantly changed phosphopeptides between EGF and mock treated cells**. (a) Quantitation results. (b) KEGG pathway. (c) Reactome pathway.

**Table S3. Pathway enrichment analysis of significantly changed phosphopeptides**. (a) Quantitation results. (b) Pathways enriched in red pulp. (c) Pathways enriched in white pulp.

**Table S4. The summary of number of identified phosphopeptides in this study and other two published works**.

**Table S5. Buffer composition for in-Tip High-pH fractionation and concatenation**.

