## Supplementary Material for "A streamlined tandem tip-based workflow for sensitive nanoscale phosphoproteomics"

Pacific Northwest National Laboratory

Richland, WA 99354

Dr. Tujin Shi

Integrative Omics Group

Biological Sciences Division

Pacific Northwest National Laboratory

Richland, WA 99354

Dr. Tao Liu

Integrative Omics Group

Biological Sciences Division

Pacific Northwest National Laboratory

Richland, WA 99354

**SI Figures:**

**Figure S1.** **The comparison of the step-by-step and integrated tip-based IMAC methods for phosphopeptides enrichment.** (a) Schematic illustration of the two methods. The number, XIC area (b), Pearson correlation, and CV (%) (c) of identified phosphopeptides purified from 10 μg of proteins from the A549 cell lysate.

**Table S5. Buffer composition for in-Tip High-pH fractionation and concatenation.**

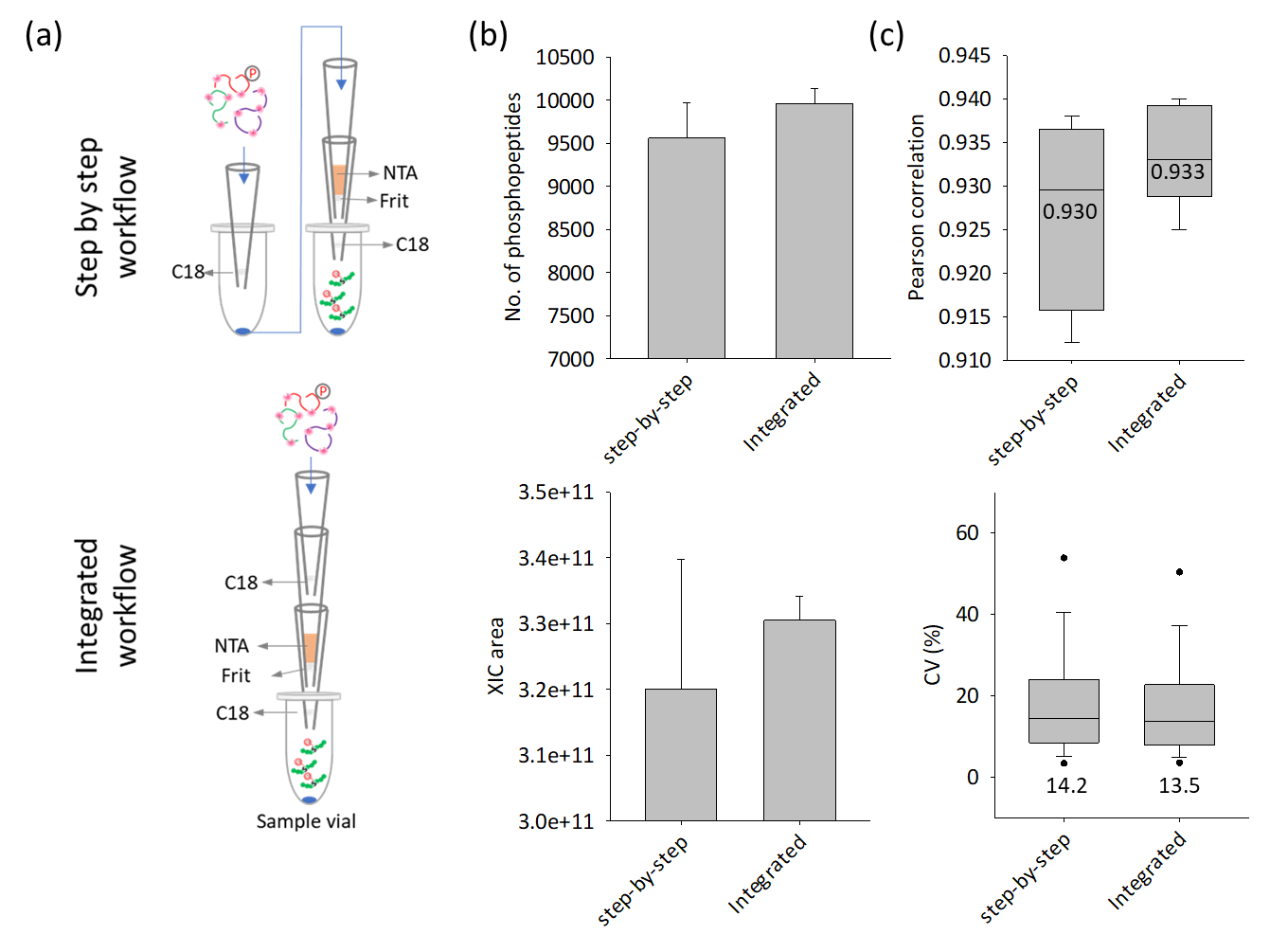

**Figure S1.** **The comparison of the step-by-step and integrated tip-based IMAC methods for phosphopeptides enrichment.** (a) Schematic illustration of the two methods. The number, XIC area (b), Pearson correlation, and CV (%) (c) of identified phosphopeptides purified from 10 μg of proteins from the A549 cell lysate.

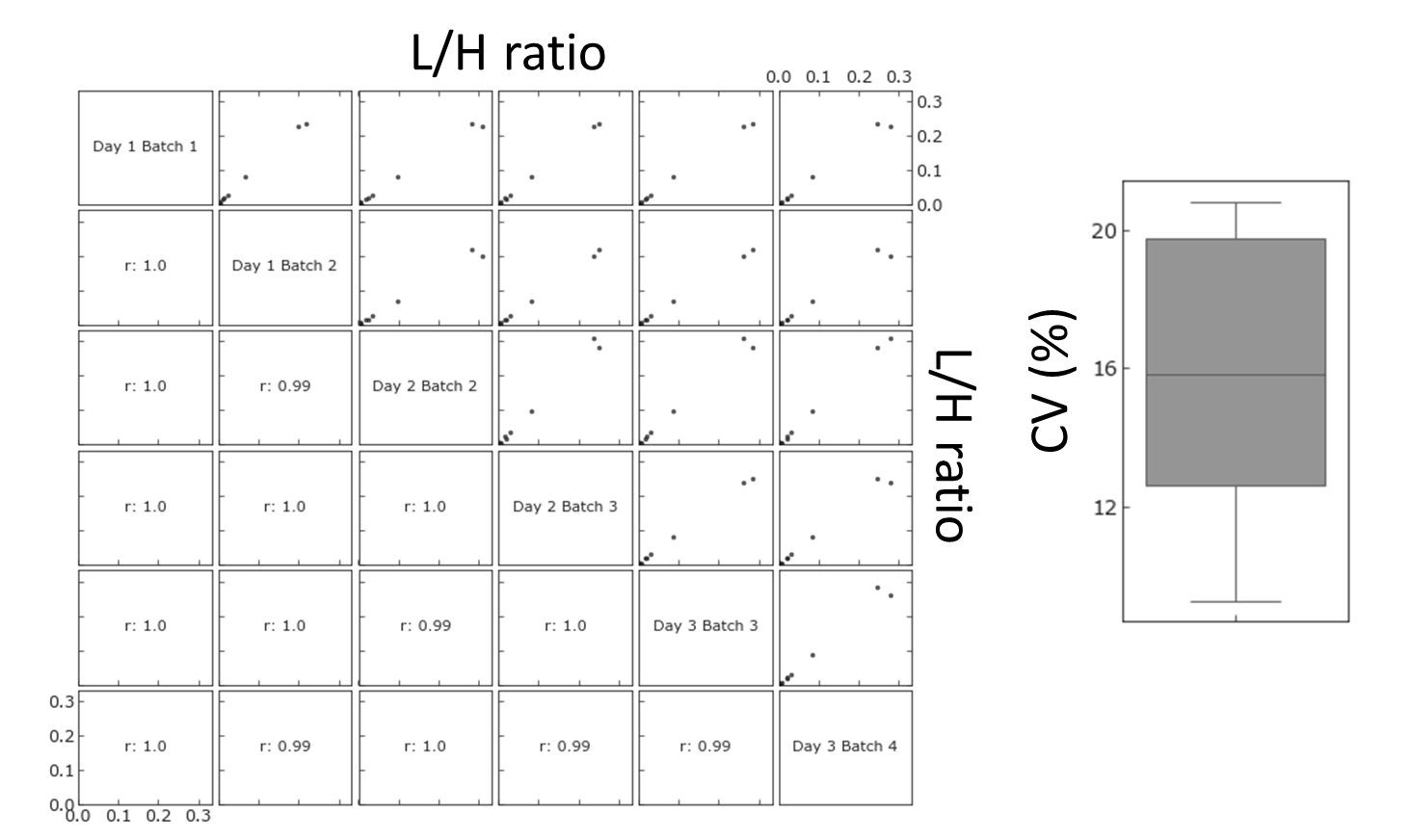

**Figure S2.** **The reproducibility of integrated tip-IMAC for phosphopeptides enrichment.** Pearson correlation and CV (%) of peak areas of 10 spiked SIL phosphopeptides (SRM data) in a reference endometrial tumor sample enriched with tip-based IMAC in different days and batches.

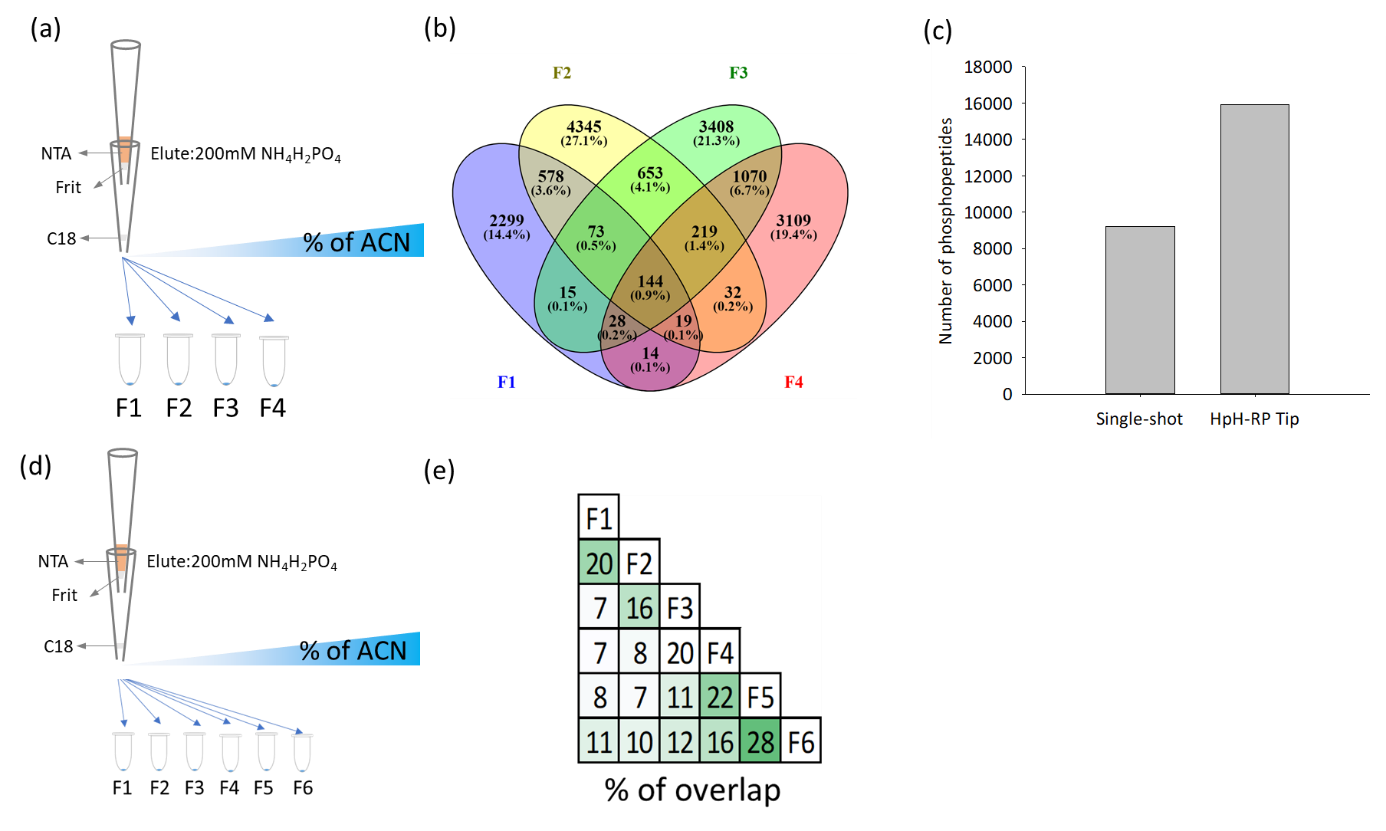

**Figure S3.** **A** **IMAC-HpH-RP tip method for microscale phosphopeptide fractionation.** (a) Using the integrated C18-IMAC-HpH-RP tip method for phosphopeptides fractionation (4 fractions for 20 μg tryptic peptides of MCF-7 cells). (b) Low overlap of the phosphopeptides identified from the 4 different fractions. (c) The number of identified phosphopeptides using the single-shot and HpH-RP tip-based fractionation method. (d) The tandem IMAC-HpH-C18 tip for phosphopeptide fractionation (6 fractions for 500 μg of tryptic peptides of MCF-7 cells). (e) Separation efficiency calculated by the percentage of common phosphopeptides between fractions.

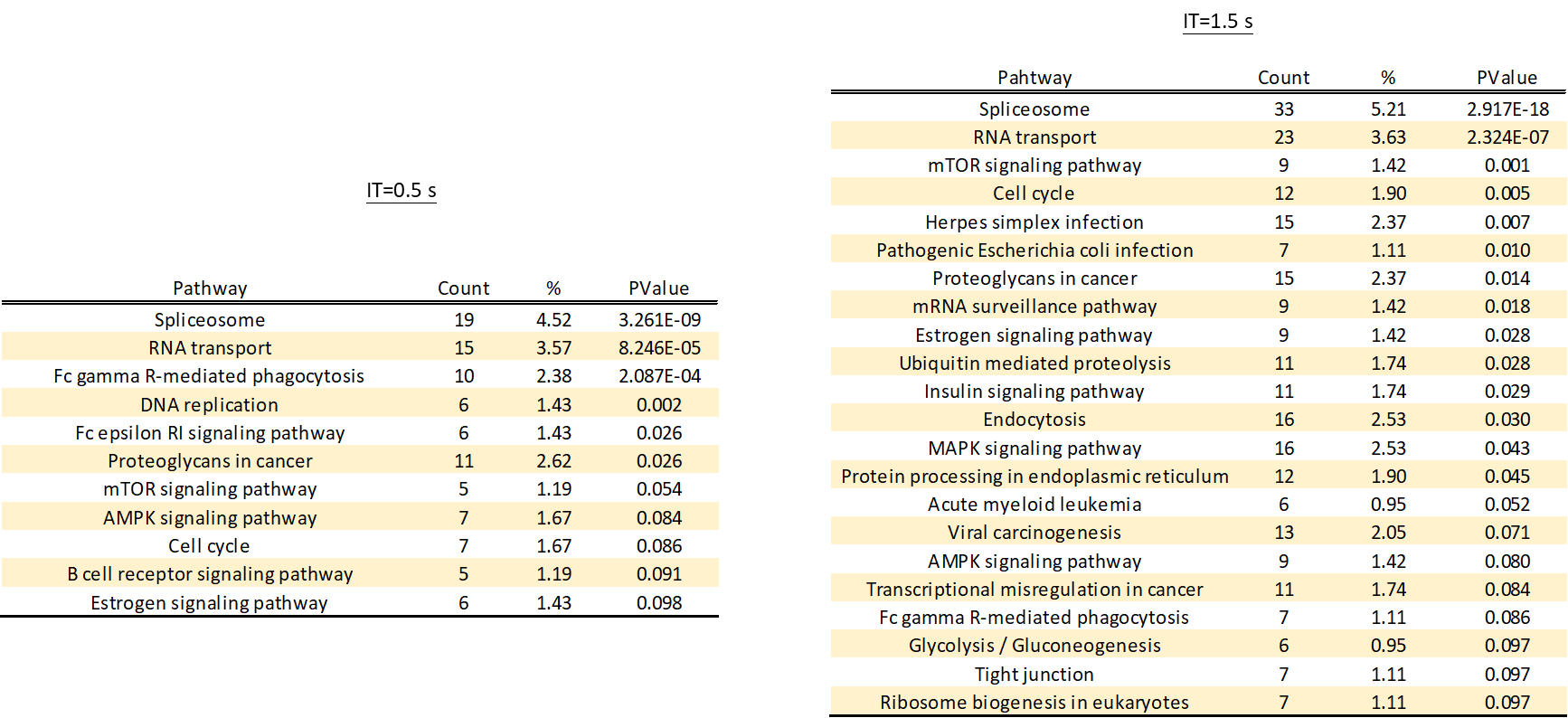

**Figure S4.** **Pathway enrichment analysis of significantly changed phosphopeptides of 10 ng tryptic digests of AML cells under different ion injection times.**

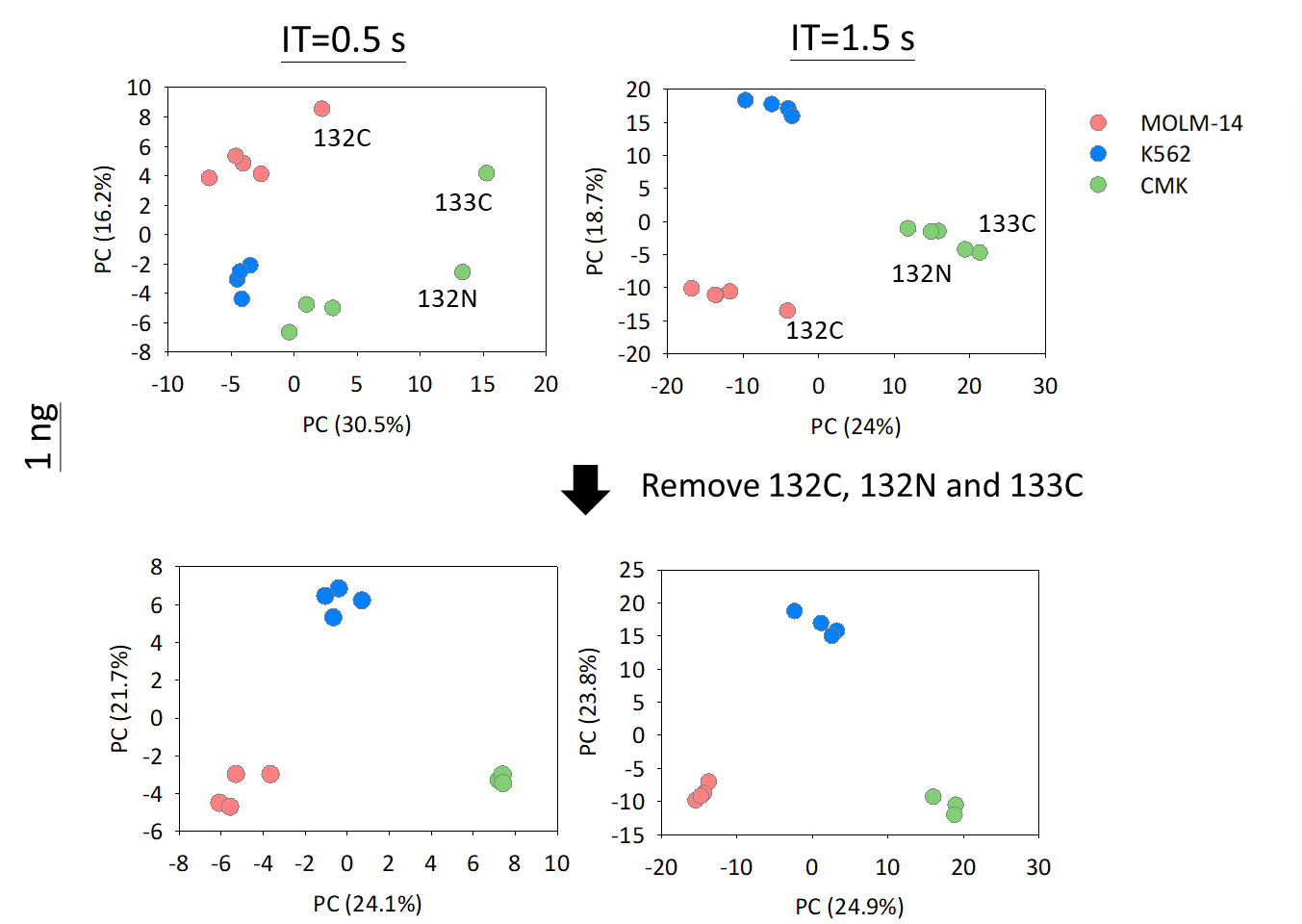

**Figure S5. Effect of ion injection time on quantitation quality in the mimic nanoscale phosphoproteome analysis**. The PCA analysis of quantified phosphopeptides of 10 ng tryptic digests of the AML cells under two different ion injection times (0.5 and 1.5 s).

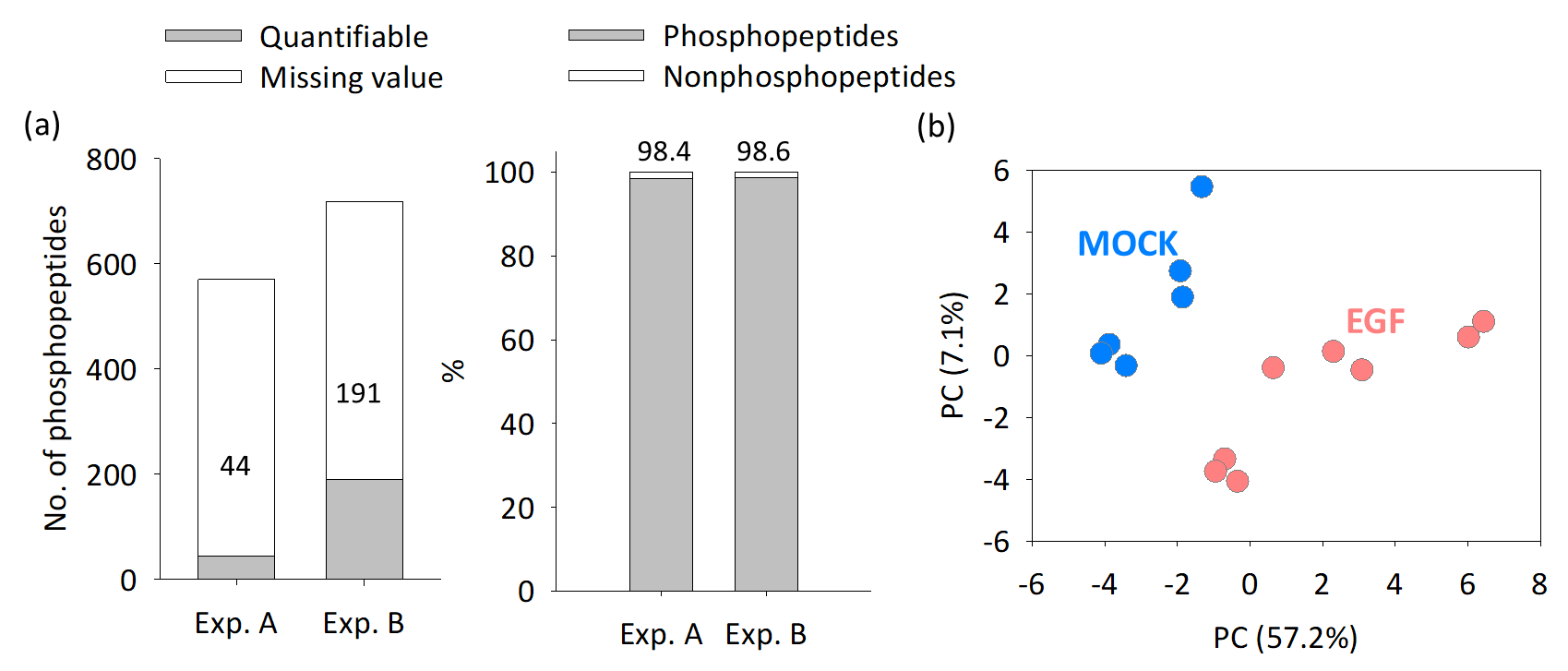

**Figure S6. Phosphoproteome analysis for 10 sorted MCF10A cells.** (a) The numbers of quantified (70% no-missing value in study samples) phosphopeptides and enrichment specificity in each TMT experiment. (b) PCA analysis shows the clustering of cells from the two different treatment conditions.

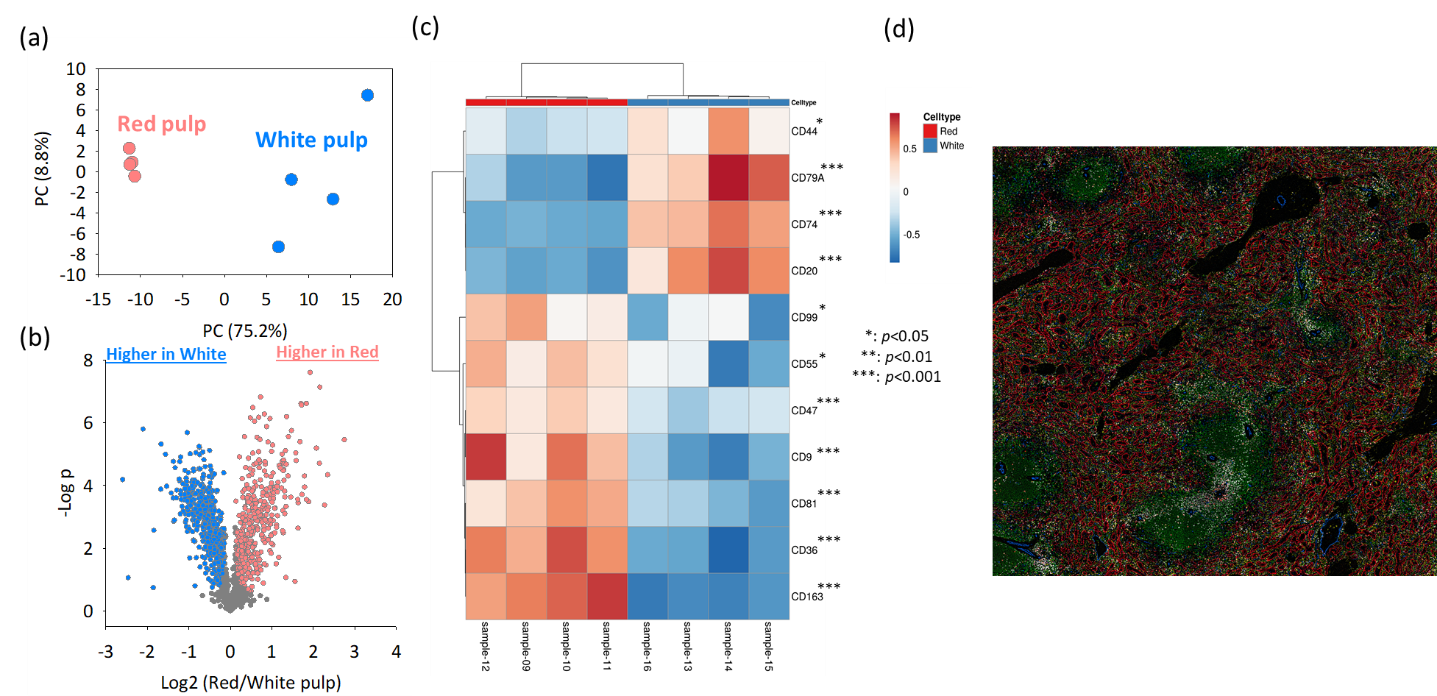

**Figure S7. Global proteome analysis of LCM-dissected human spleen tissue voxels.**  (c) PCA of the proteome data. (d) Volcano plot shows significantly changed proteins in white pulp and red pulp. (c) The altered proteins expression of surface markers. (d) Representative CODEX image of human spleen tissue: CD8α (red), CD163 (yellow), CD3e (white), CD20 (green), CD31 (blue). Scale bar, 560 µm.

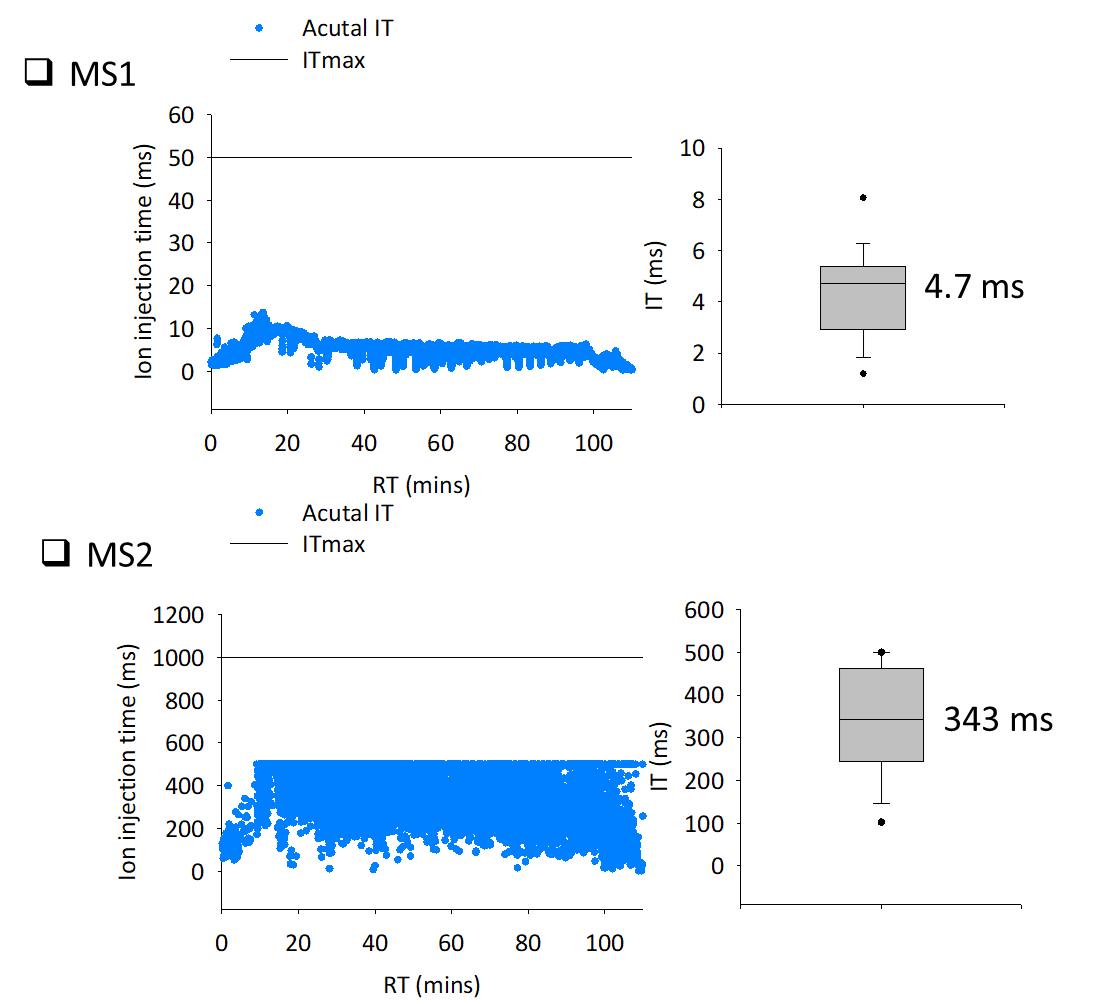

**Figure S8. The ion injection time distribution of purified phosphopeptides from 0.1** μ**g proteins from the A549 cell lysate at MS1 (a) and MS2 level.**

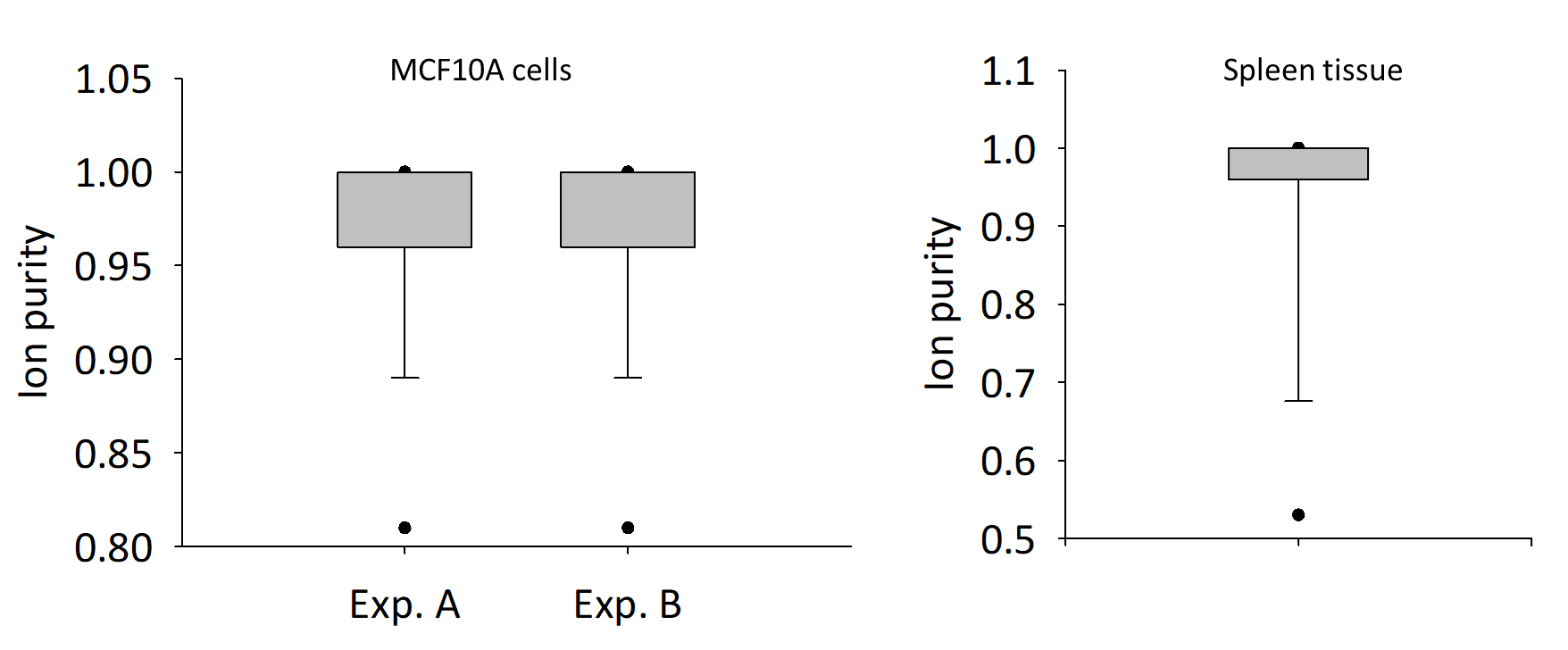

**Figure S9. The ion purity distribution of detected phosphopeptides in Figure 6 and 7.**
